# Mitotic transcription ensures ecDNA inheritance through chromosomal tethering

**DOI:** 10.1101/2025.02.12.637945

**Authors:** Ashley Nichols, Roshan Norman, Yanyang Chen, Yujin Choi, Josefine Striepen, Eralda Salataj, Eleonore Toufektchan, Richard Koche, John Maciejowski

## Abstract

Extrachromosomal DNA (ecDNA) are circular DNA bodies that play critical roles in tumor progression and treatment resistance by amplifying oncogenes across a wide range of cancer types. ecDNA lack centromeres and are thus not constrained by typical Mendelian segregation, enabling their unequal accumulation within daughter cells and associated increases in copy number. Despite intrinsic links to their oncogenic potential, the fidelity and mechanisms of ecDNA inheritance are poorly understood. Here, we show that ecDNA are protected against cytosolic mis-segregation through mitotic clustering and by tethering to the telomeric and subtelomeric regions of mitotic chromosomes. ecDNA-chromosome tethering depends on BRD4 transcriptional co-activation and mitotic transcription of the long non-coding RNA *PVT1*, which is co-amplified with *MYC* in colorectal and prostate cancer cell lines. Disruption of ecDNA-chromosome tethering through BRD4 inhibition, *PVT1* depletion, or inhibiting mitotic transcription results in cytosolic mis-segregation, ecDNA reintegration, and the formation of homogeneously staining regions (HSRs). We propose that nuclear inheritance of ecDNA is facilitated by an RNA-mediated physical tether that links ecDNA to mitotic chromosomes and thus protects against cytosolic mis-segregation and chromosomal integration.

## INTRODUCTION

Dysregulated expression of oncogenes and proto-oncogenes is a hallmark of oncogenesis. Enhanced oncogene expression often reflects copy number gains achieved through structural and numerical chromosomal aberrations^1^. Extrachromosomal DNA (ecDNA) are circular DNA molecules that capture and amplify proto-oncogenes and oncogenes in >60% of cancer types^2^. ecDNA can form when catastrophic genomic events, such as chromothripsis or breakage-fusion-bridge cycling, generate DNA fragments that escape accurate repair and/or degradation and are subsequently circularized^3–8^. Widespread detection of ecDNA across diverse cancer types, including glioblastoma, neuroblastoma, colon, breast, and others, suggest a crucial role in cancer development^2,9^. Indeed, the presence of ecDNA is associated with enhanced genetic diversity, resistance to cancer treatments, and shorter patient survival times^10^.

ecDNA possess a more open chromatin configuration that enhances accessibility to the transcriptional machinery and confers a distinct transcriptional advantage over linear oncogene amplifications^11,12^. In addition, ecDNA are tethered together by the bromodomain and extraterminal domain (BET) protein BRD4 to engage in transcriptionally active hubs that facilitate cooperative interactions across ecDNA molecules that drive oncogene expression even higher^13,14^. Combined with their high copy number presence, this favorable transcriptional profile causes ecDNA-encoded genes to be among the most highly expressed in human tumor samples^11^. Together, these qualities drive rampant transcription of oncogenes, resulting in aggressive tumor growth and treatment resistance. However, high levels of transcription place ecDNA at greater risk of transcription-replication conflicts, and the ensuing replication stress exposes a potential therapeutic vulnerability in ecDNA-containing cancers^15^.

Unlike chromosomal DNA, which segregates evenly through centromere-mediated attachment to the mitotic spindle, ecDNA lack centromeres and, therefore, segregate randomly during mitosis^16–18^. Asymmetric segregation results in an uneven distribution of ecDNA among daughter cells, leading to intratumoral and intercellular variability in oncogene copy number^19–23^. This non-Mendelian inheritance of ecDNA allows cancer cells to rapidly adapt to environmental pressures, including therapeutic interventions, by altering oncogene copy number^24^. Upon challenge with targeted therapies, ecDNA can reintegrate into non-native chromosomal loci to form chromosome anomalies termed homogeneously staining regions (HSRs)^3,25,26^. During mitosis, ecDNA remain closely associated with chromosomal DNA, thus providing a means for acentric nuclear retention^16,27^. Tethering to chromosomal DNA during cell division, or “mitotic hitchhiking”, helps maintain the presence of ecDNA in daughter cells, thus contributing to the persistence of ecDNA-driven oncogene amplification in cancer. However, ecDNA have also frequently been observed in nuclear atypia, such as micronuclei, which occur as a result of mitotic error^3,28,29^. The frequency and fidelity of ecDNA-chromosome tethering are thus poorly understood.

Here, we report that ecDNA achieve nuclear inheritance by tethering to the telomeric and subtelomeric regions of mitotic chromosomes. By focusing our analysis on a subset of *MYC*-amplifying, ecDNA-containing, colorectal, and prostate cancer cell lines, we discovered that the long non-coding RNA *PVT1*, which is frequently co-amplified with *MYC*, is required to promote ecDNA nuclear inheritance. Furthermore, we find that ecDNA bypass normal blocks against mitotic transcription and that inhibition of mitotic transcription or depletion of *PVT1*, a long non-coding RNA often co-amplified with *MYC* on ecDNA, results in cytosolic mis-segregation, ecDNA chromosomal reintegration, and HSR formation. Our results suggest that active mitotic transcription of ecDNA-encoded genes promotes attachment to mitotic chromosomes and thus safeguards ecDNA nuclear inheritance.

## RESULTS

### ecDNA-chromosome tethering preserves nuclear inheritance

To study ecDNA segregation dynamics, we used fluorescence *in situ* hybridization (FISH) probes to track ecDNA segregation in a panel of ecDNA-containing human cancer cell lines (Figure 1A). The panel of cell lines included representatives from colorectal cancer (COLO320-DM; *MYC*+ ecDNA), glioblastoma (GBM39; *EGFR*+ ecDNA), gastric cancer (SNU16; *FGFR2+* and *MYC*+ ecDNAs), prostate cancer (PC3; *MYC*+ ecDNA), and a cervical cancer line in which ecDNA were generated in response to methotrexate treatment (HeLa-DM; methotrexate-induced *DHFR*+ ecDNA^3^). In several instances, experiments included spontaneously occurring clonal derivatives in which high copy ecDNA reintegrated into chromosomes to form HSRs (COLO320-HSR, GBM39-HSR, HeLa-HSR). Relative FISH signals across daughter cells were used to measure the symmetry of ecDNA segregation, while the proportion of nuclear to total cellular FISH signals was used to assay nuclear segregation fidelity. Anti-Aurora B immunofluorescence (IF) identified recently divided daughters. FISH measurements validated expected observations of asymmetric ecDNA accumulation across daughters (Figure S1A)^18,22,23^ and revealed that ecDNA localize mainly to the nucleus (Figures 1A and 1B). Quantification of the proportion of nuclear to total FISH signal confirmed that ecDNA localization is primarily restricted to the nucleus, albeit at a lower fidelity than chromosomally integrated HSRs (Figure 1B). ecDNA nuclear localization did not correlate with the total FISH signal, indicating that nuclear segregation fidelity is not impacted by ecDNA copy number (Figure S1B).

**Figure 1.**
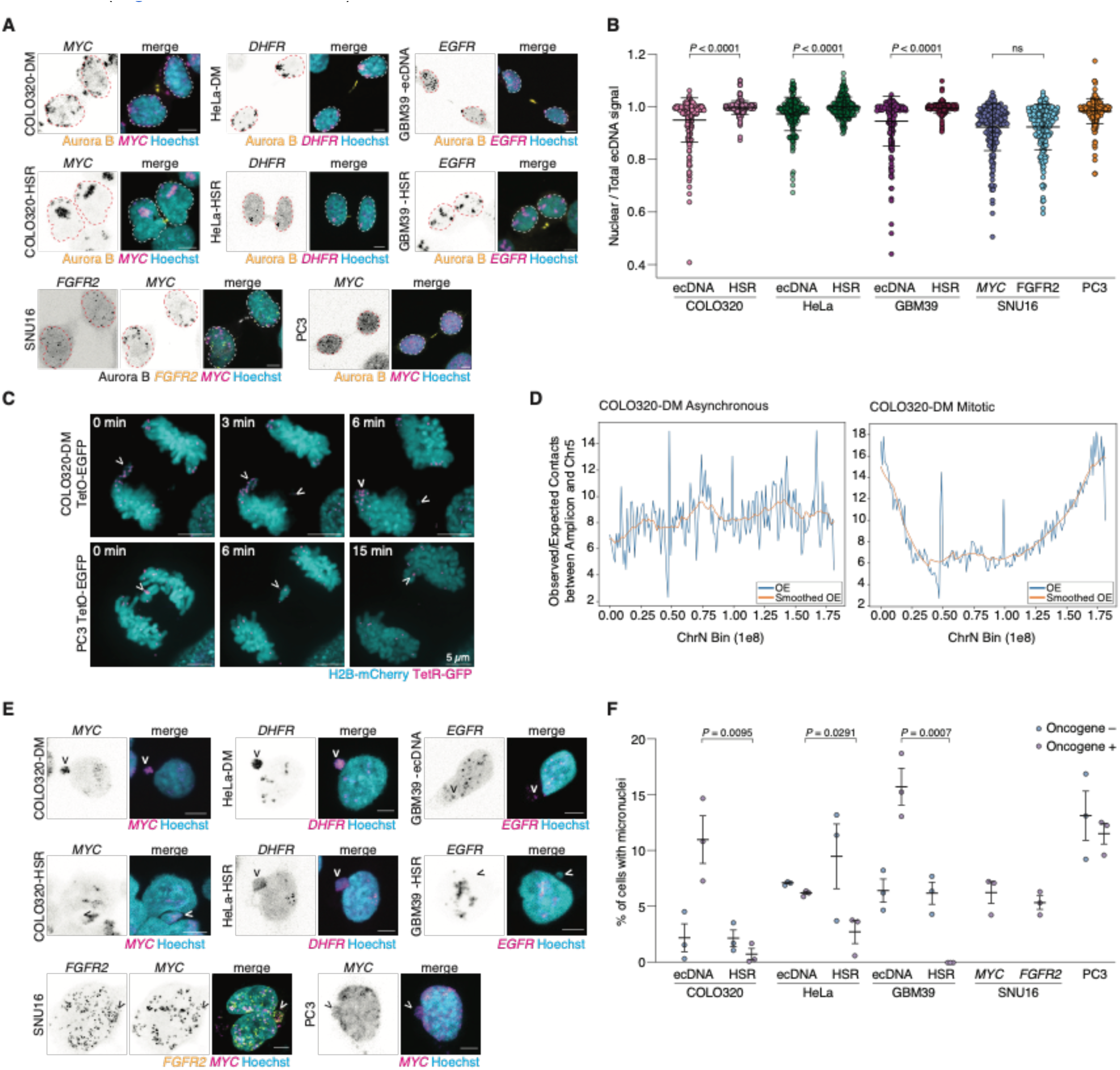
ecDNA achieve high fidelity nuclear inheritance by tethering to mitotic chromosomes. (A) Immunofluorescence-DNA FISH of newly formed daughter cells in ecDNA-positive or HSR-positive cancer cell lines. Recently divided daughter cell pairs were identified by anti-Aurora B immunofluorescence and gene amplifications were quantified with sequence-specific FISH probes. Nuclei borders are indicated. (B) Quantification of ecDNA nuclear segregation fidelity in newly formed daughter cells as shown in (A). Mean±s.d. of *n* = 3 experiments (100 pairs of daughter nuclei, >25 pairs analyzed per experiment) are shown. (C) Live-cell imaging of COLO320-DM TetO-EGFP and PC3 TetO-EGFP cells expressing H2B-mCherry and TetR-GFP. Arrows mark ecDNAs segregating into daughter nuclei or cytosol. Insets mark time in minutes since anaphase (t = 0). (D) Linear chromosome map demonstrating the log10 fold-change of observed over expected (OE) valid contacts between the COLO320-DM amplicon and chromosome 5 in asynchronous versus mitotic samples. All valid pairs were binned into 1 Mb bins and plotted. (E) Identification of oncogene-positive or oncogene-negative micronuclei in cancer cell line panel through oncogene FISH using sequence-specific FISH probes. Arrows indicate micronuclei. (F) Quantification of the percentage of cells containing at least one oncogene-positive or oncogene-negative micronucleus as in (E). Mean±s.d. of *n* = 3 experiments shown (>100 primary nuclei analyzed per experiment). *P* values were calculated via unpaired *t*-tests. Scale bars = 5 μm. See also Figure S1-S3.

We reasoned that the restricted nuclear localization of ecDNA could reflect an ecDNA nuclear segregation mechanism or degradation of ecDNA mis-segregated into the cytosol. To distinguish between these possibilities, we used super-resolution, live-cell imaging to track ecDNA dynamics during mitosis. ecDNA were visualized through the insertion of a Tet-operator (TetO) array into *MYC*+ ecDNAs in COLO320-DM and PC3 cells and labeling with TetR-eGFP^13,22^. In agreement with our IF-FISH results, live-cell imaging revealed that ecDNA rarely mis-segregate into the cytosol and display a high fidelity for nuclear inheritance (Figure 1C; Video S1, S2). ecDNA organized into clusters that are associated with or “hitchhiked” on mitotic chromosomes. ecDNA clusters frequently segregated as one entity, thus contributing to their ability to rapidly amplify associated oncogenes. ecDNA appeared to often make contact with telomeric or subtelomeric regions of mitotic chromosomes (Figure S1C). In several instances, we could observe groups of ecDNA lag and likely mis-segregate as a large cluster after failing to attach to a mitotic chromosome (Figure S1D). These data indicate that ecDNA segregation is an active process, consistent with observations from decades ago^16,30–33^ that ecDNA-chromosome hitchhiking facilitates ecDNA nuclear segregation.

To investigate ecDNA-chromosome contacts further, we performed genome-wide Hi-C sequencing of asynchronous and mitotically arrested COLO320-DM cells. Mitotically arrested cells were prepared through FACS-based isolation of nocodazole-treated cells (Figures S2A-C; Methods). Importantly, live-cell imaging confirmed that ecDNA-chromosome contacts were preserved during nocodazole arrest (Figure S2A). Hi-C sequencing showed that ecDNA made widespread contacts across chromosomes in asynchronous cells. In contrast, ecDNA-chromosome contacts underwent a dramatic rearrangement in mitotic cells, which showed enrichment of contact sites at chromosome ends (Figure 1D; Figure S2D and S2E). ecDNA contacts with chromosome ends were widespread throughout the genome. Based on these data, we conclude that ecDNA achieve high fidelity nuclear inheritance by tethering to the telomeric and subtelomeric regions of mitotic chromosomes. This high fidelity for nuclear segregation exhibited by ecDNA is distinct from linear, acentric chromosome fragments produced by DNA double-strand breaks or centromere dysfunction, which frequently mis-segregate into the cytosol^7,34^.

Although the majority of ecDNA were segregated into the nucleus, further analysis of cellular ecDNA distribution revealed that ecDNA could be detected within micronuclei (Figures 1E and 1F). ecDNA-containing micronuclei appeared to be composed of multiple ecDNA copies, suggesting that ecDNA may mis-segregate as a large, clustered unit. Micronucleation is associated with DNA damage and catastrophic chromosome rearrangement^35–38^. We, therefore, assessed DNA damage in ecDNA-containing micronuclei using immunofluorescence against the DNA damage marker γH2AX. As expected, ecDNA+ micronuclei, marked by oncogene DNA FISH, exhibited high levels of γH2AX, indicating extensive DNA damage (Figure S3A and S3B). Chromosomes isolated in micronuclei are transcriptionally silenced^39^. Analysis of active transcription using nascent *MYC* RNA-FISH probes showed that ecDNA exhibit similar transcriptional silencing within micronuclei (Figure S3C and S3D).

Together, these results indicate that ecDNA micronucleation is associated with extensive DNA damage and transcriptional silencing of ecDNA-encoded genes, such as *MYC*, and thus highlight the risks of cytosolic mis-segregation.

### ecDNA mitotic clustering, chromosome tethering, and maintenance depend on BRD4

Several proteins or protein complexes have previously been implicated in the clustering or crosslinking of chromosomes or acentric chromosome fragments during mitosis^40–42^. These include the CIP2A-TOPBP1 complex, which associates with DNA lesions generated at micronuclei to cluster pulverized chromosomes together upon mitotic entry, and the small DNA binding protein BAF, which bridges distant DNA sites to shape nuclear assembly at mitotic exit^40–42^. To determine whether these proteins guide ecDNA segregation, we used super-resolution, live-cell imaging to track ecDNA segregation dynamics in *CIP2A* and *BAF*-depleted cells (Figure S4A-D). Neither *CIP2A* nor *BAF* depletion affected ecDNA clustering, ecDNA-chromosome tethering, or ecDNA nuclear segregation, indicating that these proteins are dispensable for guiding ecDNA segregation (Figure S4A-D). Likewise, inhibition of the spindle assembly checkpoint by treatment with Mps1 inhibitors failed to impact ecDNA nuclear segregation (Figure S4E and S4F).

Mitotic ecDNA clustering was reminiscent of the prior reports of ecDNA hubs, a phenomenon in which clusters of 10-100 ecDNAs form during interphase to drive intermolecular enhancer-gene interactions and further elevate oncogene expression^13^. Interphase ecDNA clustering depends on the BET protein BRD4^13^. To test if mitotic ecDNA dynamics are similarly dependent on BRD4, we treated cells with the BET inhibitor JQ1^43^ and tracked ecDNA segregation in recently divided COLO320-DM cells by IF-FISH (Figures 2A and 2B). JQ1 treatment significantly decreased total nuclear ecDNA signal as measured by *MYC* DNA FISH and increased the rate of cytosolic mis-segregation, often in the form of large micronuclei. Live-cell imaging further revealed that JQ1 treatment dispersed mitotic ecDNA clusters and increased cytosolic mis-segregation without affecting chromosome segregation dynamics in COLO320-HSR control cells (Figures 2C and 2D; Figure S4G and S4H; Videos S3-S4). Degradation of endogenous BRD4 by treating BRD4-HaloTag knock-in COLO320-DM cells with HaloPROTAC led to similar increases in ecDNA mis-segregation suggesting that off-target effects of JQ1 were not responsible for the observed phenotypes (Figures 2C, 2D; Figure S4I). JQ1 treatment elicited similar effects on ecDNA segregation in PC3 cells, indicating that these results extend to other *MYC*-amplifying ecDNA species (Figures 2E and 2F). Together, these data indicate that BRD4 is necessary for mitotic clustering and nuclear segregation of *MYC-* amplifying ecDNA.

**Figure 2.**
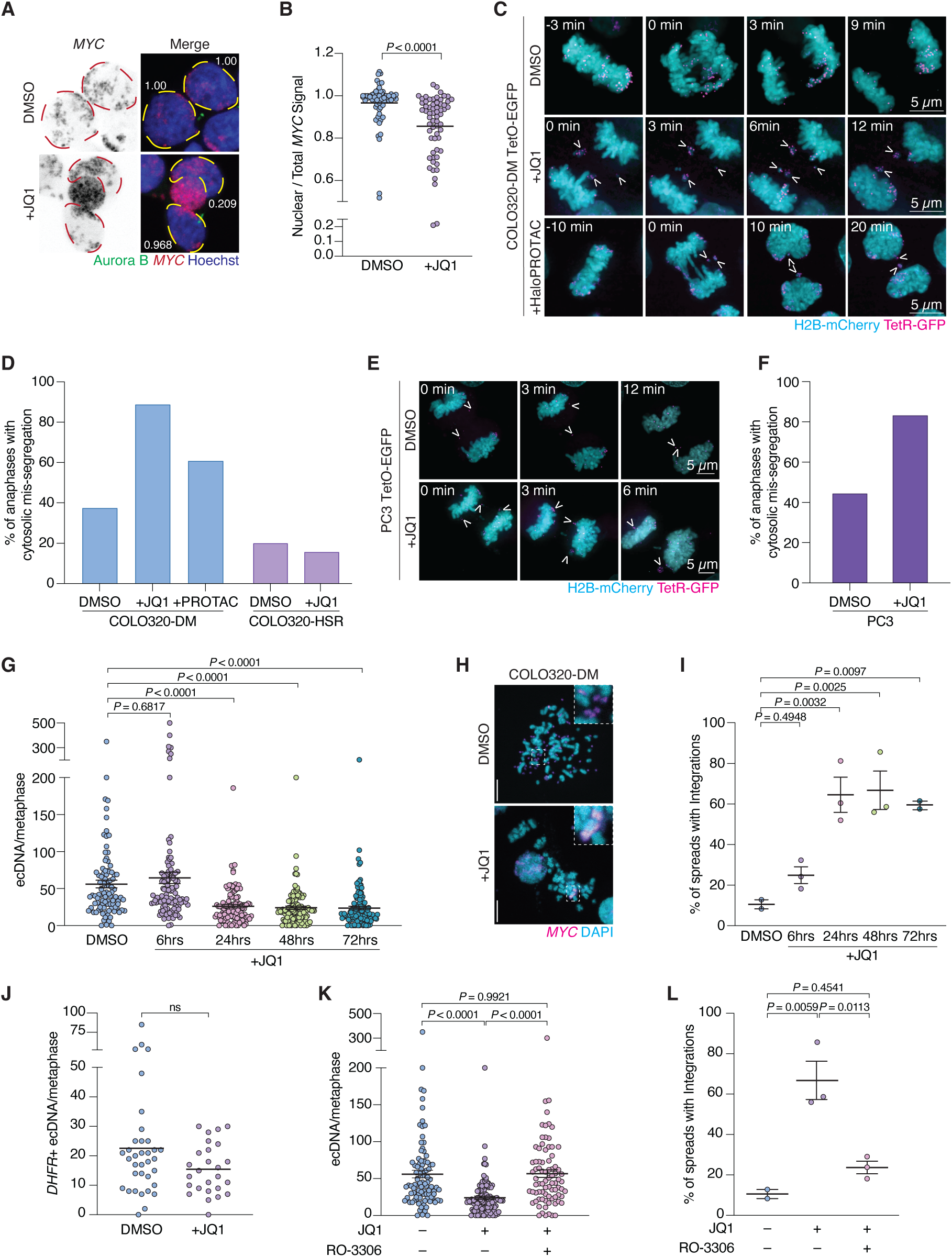
BRD4 safeguards ecDNA nuclear segregation and ecDNA maintenance. (A) Immunofluorescence-DNA FISH in COLO320-DM cells treated with DMSO or JQ1 for 6 hours. Aurora B staining identifies recently divided daughter cell pairs, and ecDNAs are labeled with a *MYC* specific DNA FISH probe. Primary nuclei borders are highlighted. Numerical labels indicate ecDNA nuclear fidelity associated with each daughter cell. (B) Quantification of ecDNA nuclear localization in COLO320-DM cells treated with DMSO or JQ1 as shown in (A). *n* = 1 experiment (>30 daughter cell pairs per replicate). *P* value was calculated with an unpaired *t*-test. (C) Live-cell imaging of COLO320-DM TetO-EGFP cells expressing H2B-mCherry and TetR-GFP. Arrows mark mis-segregating ecDNA. Labels mark time in minutes since anaphase (t = 0). (D) Quantification of ecDNA mis-segregation into the cytosol as in (C) and Figure S4H. (COLO320-DM: DMSO *n* = 16 cells, JQ1 *n* = 18 cells, HaloPROTAC *n* = 23; COLO320-HSR: DMSO *n* = 20 cells, JQ1 *n* = 19 cells). (E) Live-cell imaging of PC3 TetO-EGFP expressing H2B-mCherry and TetR-GFP. Arrows mark mis-segregating ecDNA. Insets mark time in minutes since anaphase (t = 0). (F) Quantification of live-cell imaging from (E) showing percentage of anaphases demonstrating ecDNA cytosolic mis-segregation (DMSO *n* = 18; JQ1 *n* = 12). (G) ecDNA counts from metaphase spreads of COLO320-DM cells treated with DMSO or JQ1 for indicated time periods. Mean±s.e.m. of *n* = 3 experiments (100 metaphase spreads total, >25 metaphase spreads analyzed per experiment) are shown. (H) *MYC* FISH in COLO320-DM spreads treated with DMSO or JQ1 for 48 hours. Insets display examples of *MYC* amplification on ecDNA or as HSR. (I) Quantification of metaphase DNA FISH showing average number of metaphase spreads depicting chromosomal integration of *MYC* ecDNA in COLO320-DM cells treated with JQ1. Mean±s.d. of *n* = 2-3 experiments (>25 spreads analyzed per experiment) are shown. (J) Quantification of ecDNA in HeLa-DM cells treated with DMSO or JQ1. Mean of *n* = 1 experiments (>25 spreads per condition) are shown. (K) Quantification of ecDNA number in metaphase spreads from COLO320-DM cells treated as indicated. Mean±s.e.m. of *n* = 2-3 experiments (>25 spreads analyzed per replicate) are shown. (L) Quantification of ecDNA chromosomal integration in COLO320-DM cells treated with JQ1 and RO-3306. Mean±s.d. of *n* = 2-3 experiments (>25 spreads analyzed per replicate) are shown. Unless otherwise noted *P* values were calculated by one-way ANOVA with Tukey’s multiple comparisons test. ns = not significant. Scale bars = 5 μm. See also Figure S4 and S5.

Next, we assessed how JQ1 treatment affects ecDNA maintenance. Analysis of metaphase spreads showed that prolonged JQ1 treatment (>24hrs) reduced ecDNA number (26.2±24.1 ecDNA after 24hr JQ1 vs 56.1±50.2 s.d. ecDNA in DMSO controls) (Figure 2G; Figure S5A). BRD4 hypomorph subclones generated through CRISPR-Cas9 editing showed similar reductions in ecDNA number (Figure S5B and S5C). Surprisingly, prolonged JQ1 treatment did not elicit corresponding reductions in *MYC* expression levels despite decreased ecDNA number (Figure S5D and S5E). Consistently, shallow WGS indicated that the reductions in ecDNA number seen following prolonged JQ1 treatment did not lead to corresponding decreases in the copy number of the *MYC* locus or nearby regions that are captured and co-amplified on ecDNA (Figure S5F). Conservation in copy number was explained by increased ecDNA chromosomal reintegration and HSR formation (Figures 2H and 2I; Figure S5G). HSR formation was rapid, occurring within 24h of JQ1 treatment. In contrast to results in COLO320-DM cells, JQ1 treatment did not reduce ecDNA number or promote chromosomal reintegration of *DHFR*-amplifying ecDNA in HeLa-DM cells (Figures 2J; Figure S5H and S5I). Co-treatment with the CDK1 inhibitor RO-3306 and JQ1 preserved ecDNA number and blocked HSR formation, indicating that mitotic progression is a critical intermediate in these processes (Figures 2K and 2L). Together, these data indicate that BRD4 promotes ecDNA nuclear inheritance, which protects against chromosomal reintegration, and support a model in which HSR formation can occur through the simultaneous integration of multiple ecDNA copies^44^.

### BRD4 enhances ecDNA-chromosome tethering by promoting ecDNA transcription

BRD4 is enriched within interphase ecDNA hubs where it is proposed to physically link *MYC*-amplifying ecDNA molecules by participating in ecDNA-ecDNA *trans* interactions^13^. Based on these observations we examined BRD4 mitotic localization to test whether it promotes ecDNA mitotic clustering and chromosome tethering by engaging in a similar physical linkage. However, IF-FISH using anti-BRD4 antibodies and live-cell imaging of endogenously tagged HaloTag-BRD4^13^ failed to identify a significant overlap between BRD4 and ecDNA on mitotic chromosomes, indicating that BRD4 likely plays a more indirect role in these processes (Figure S6A).

Interphase ecDNA hubs depend on ecDNA-borne nascent RNAs, such as the long non-coding RNA *PVT1* that is often co-amplified with *MYC* on ecDNA^45^. *PVT1* depletion scatters interphase ecDNA hubs, disrupts ecDNA-associated trans-activation, and decreases oncogene expression^45^. We, therefore, reasoned that mitotic ecDNA clustering and segregation may similarly depend on *PVT1*. Indeed, live-cell imaging revealed that *PVT1* depletion using antisense oligonucleotides (ASOs) drove ecDNA mis-segregation into the cytosol in COLO320-DM and PC3 cells (Figures 3A-D; Figure S6B and S6C). As expected, *PVT1* depletion did not perturb chromosome segregation in COLO320-HSR control cells (Figure 3B; Figure S6D). Of further similarity to JQ1 treatment, *PVT1* knockdown by shRNA infection decreased ecDNA number while increasing the frequency of ecDNA chromosomal re-integration and HSR formation (Figures 3E-G; Figure S6E and S6F). Together, these observations indicate that the ecDNA-encoded *PVT1* RNA plays a critical role in ecDNA-chromosome tethering and ecDNA segregation. Interestingly, JQ1 treatment significantly diminished *PVT1* expression levels (Figure S6G), likely by interfering with the pivotal roles of BRD4 in transcription^46^. Thus, the impact of BRD4 inhibition on ecDNA segregation and interphase hub formation may, at least partially, be explained by *PVT1* depletion.

**Figure 3.**
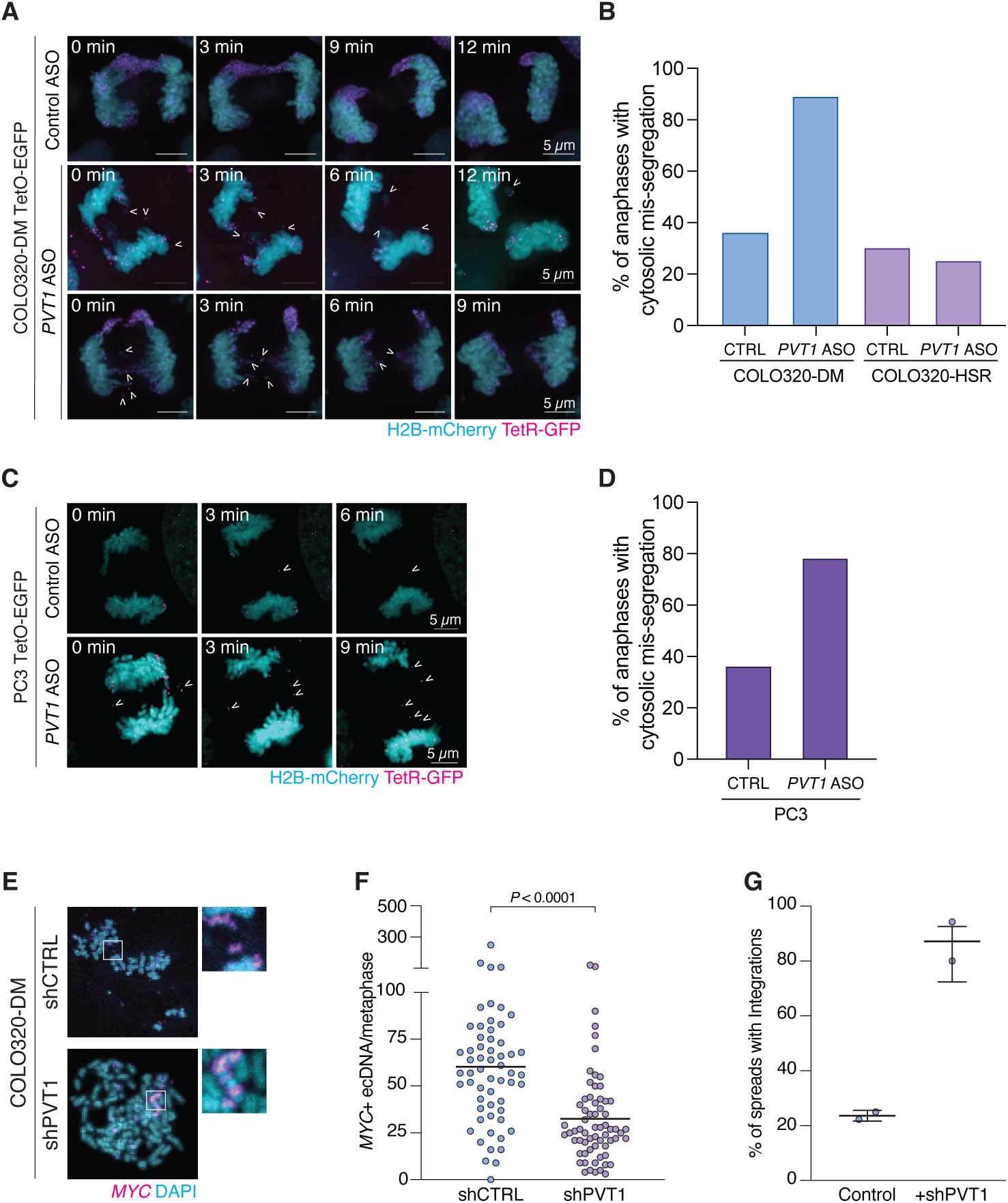
*PVT1* depletion induces ecDNA mis-segregation and chromosomal integration. (A) Live-cell imaging of COLO320-DM TetO-EGFP cells expressing H2B-mCherry and TetR-GFP transfected with control or *PVT1-*specific ASO. Arrows label ecDNAs mis-segregating into the cytosol. (B) Quantification percentage of cells demonstrating ecDNA cytosolic mis-segregation upon PVT1 depletion as shown in (A). (COLO320-DM: Control *n* = 25, *PVT1* ASO *n* = 28; COLO320-HSR Control and *PVT1* ASO *n* = 20 each). (C) Live-cell imaging of PC3 TetO-EGFP expressing H2B-mCherry and TetR-GFP after transfection with indicated ASOs. Arrows mark tagged ecDNAs demonstrating cytosolic mis-segregation. (D) Quantification of anaphases displaying cytosolic mis-segregation as depicted in (C) in PC3 TetO-EGFP cells (Control *n* = 14, *PVT1* ASO *n* = 18). (E) Representative metaphase *MYC* FISH in COLO320-DM cells overexpressing shCTRL or shPVT1. Insets depict ecDNA-amplified *MYC* versus *MYC* integration in indicated cell lines. (F) Quantification of the number of ecDNA in metaphase spreads from COLO320-DM cells expressing shCTRL or shPVT1. Mean of *n* = 2 experiments (>30 metaphase spreads per experiment) are shown. *P* values were calculated by unpaired *t*-tests. (G) Percentage of COLO320-DM spreads containing *MYC* integrations after expression of shCTRL or shPVT1 as shown in (E, F). Mean±s.d. of *n* = 2 experiments (>30 metaphase spreads per experiment) are shown. Scale bars = 5 μm. Insets in live-cell imaging mark time in minutes since anaphase (t = 0). See also Figure S6.

### Mitotic ecDNA transcription promotes nuclear inheritance

Unlike *GAPDH* transcripts, which disperse throughout the nucleoplasm during mitosis, RNA FISH revealed that *PVT1* transcripts remain associated with ecDNA throughout mitosis, suggesting that *PVT1* may play an active intra-mitotic role in ecDNA-chromosomal tethering (Figure S7A). Indeed, favorable transcriptional profiles and resulting pervasive transcription raised the possibility that ecDNA may resist global mitotic repression to remain transcriptionally active during mitosis^47–49^. To test this directly, we used intron-directed *PVT1* RNA FISH probes to monitor nascent transcription in interphase and mitotic COLO320-DM cells (Figures 4A-C). Since most introns are removed co-transcriptionally and quickly degraded, intronic RNA FISH probes are commonly used tools to measure nascent transcription^50,51^. In agreement with recently reported results^18^, these experiments showed that nascent *PVT1* transcripts could be detected throughout mitosis (Figure 4A). Quantification of nascent *PVT1* RNA signal in mitosis demonstrated a small but significant decrease relative to interphase cells (Figure 4B). Treatment with triptolide, which prevents the formation of “transcription bubbles” and thus blocks transcription initiation^18,52^, decreased nascent *PVT1* FISH signal in interphase and mitotic cells, thus highlighting the specificity of this assay for active transcription (Figures 4B and 4C; Figure S7B). As expected, *PVT1* intronic and exonic probes were sensitive to RNase A treatment (Figure S7C). Intronic *PVT1* RNA FISH signal could be detected in mitotic COLO320-HSR cells, indicating that these anomalous chromosomal structures may also possess an aberrant, mitotic transcriptional activity (Figure S7D). Based on these experiments, we conclude that *MYC*-amplifying ecDNA species resist global, repressive blocks on transcription throughout mitosis to actively transcribe genes like *PVT1*.

**Figure 4.**
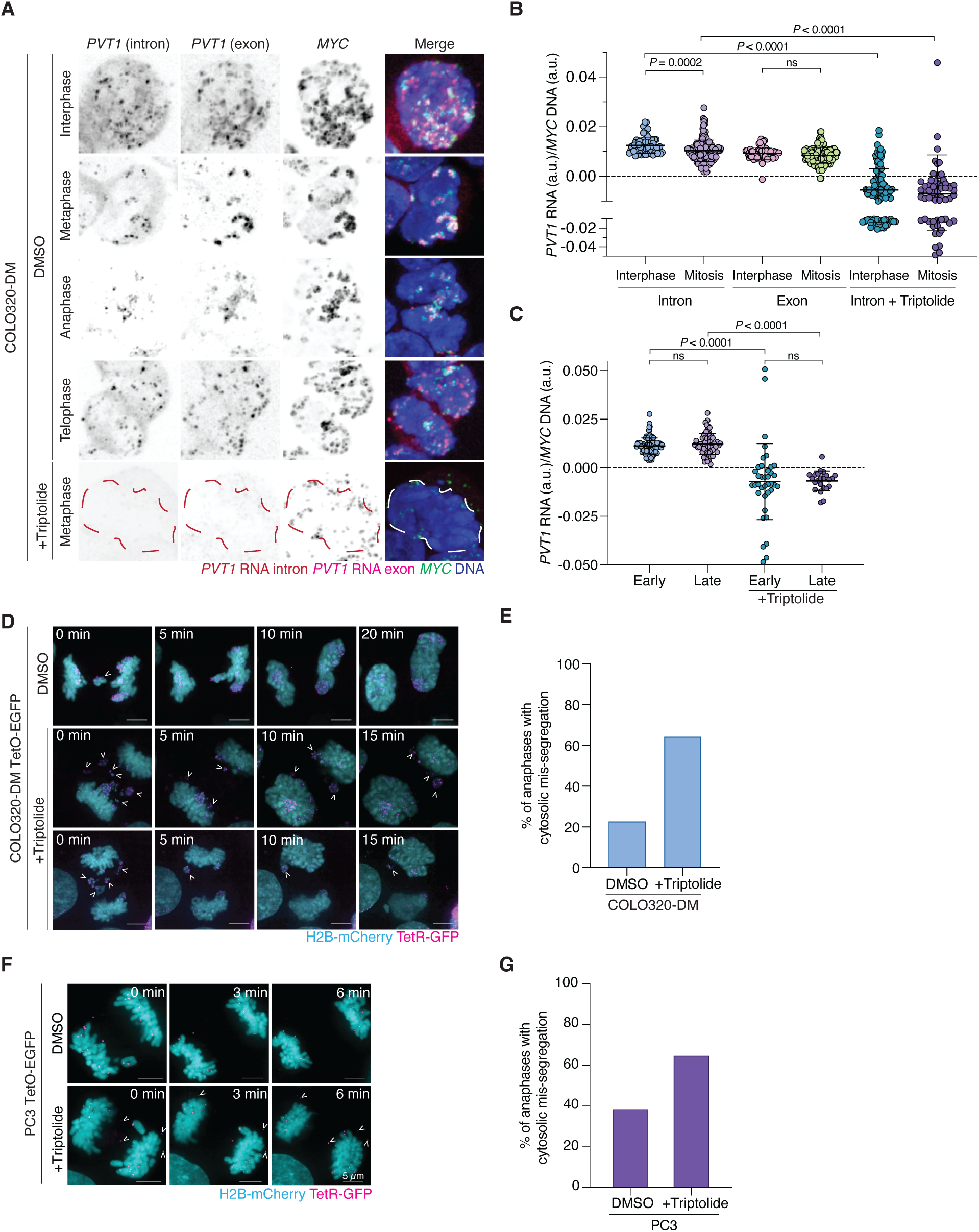
Active mitotic transcription of ecDNA enables nuclear segregation. (A) Combined RNA/DNA FISH in COLO320-DM cells treated with DMSO or triptolide during various stages of mitosis or interphase. *PVT1* transcripts were visualized with an intron and an exon probe, and ecDNA was stained with a *MYC* DNA FISH probe. Nucleus border is highlighted in the triptolide-treated, representative image. (B) Quantification of combined RNA/DNA FISH as in (A). Signal intensity of indicated PVT1 RNA probe at ecDNA+ regions normalized to *MYC* ecDNA signal intensity. Mean±s.d. of *n* = 1 experiment (>30 interphase and mitotic nuclei quantified with between 1-6 ecDNA+ regions measured per cell) are shown. (C) Quantification of combined RNA/DNA FISH in mitotic COLO320-DM cells from (A,B) stratified by phase of mitosis. Mean±s.d. of *n* = 1 experiment (>30 interphase and mitotic nuclei quantified with between 1-6 ecDNA+ regions measured per cell) are shown. Early mitosis indicates cells in prophase, prometaphase, and metaphase, while late mitosis indicates cells fixed in anaphase and telophase. (D) Live-cell imaging in COLO320-DM TetO-EGFP cells expressing H2B-mCherry and TetR-GFP treated with DMSO or triptolide. Arrows mark ecDNA demonstrating cytosolic mis-segregation. (E) Quantification of live-cell imaging in (D) showing percentage of cells with mis-segregating ecDNA (DMSO *n* = 22, Triptolide *n* = 28). (F) Live-cell imaging of PC3 TetO-EGFP cells expressing H2B-mCherry and TetR-GFP treated with DMSO or triptolide. Arrows indicate mis-segregating ecDNA. (G) Quantification of anaphases from live-cell imaging in (F) demonstrating ecDNA mis-segregation (DMSO *n* = 13, Triptolide *n* = 17). *P* values were calculated using unpaired *t*-tests. Scale bars = 5 μm Insets in live-cell imaging indicate time since anaphase onset (t = 0). ns = not significant. See also Figure S7.

Next, to investigate potential functional impacts of mitotic ecDNA transcription on ecDNA segregation, we performed super-resolution, live-cell imaging of triptolide-treated COLO320-DM cells. Similar to images generated following JQ1 treatment or *PVT1* depletion, these experiments showed that a brief triptolide treatment of just two hours was sufficient to disrupt ecDNA clustering and ecDNA-chromosome tethering and drive cytosolic mis-segregation (Figures 4D and 4E; Video S5). Triptolide treatment similarly interfered with ecDNA clustering and nuclear segregation in mitotic PC3 cells (Figures 4F and 4G). In contrast, triptolide treatment failed to elicit significant impacts on chromosome segregation in COLO320-HSR control cells (Figure S7E and S7F). Taken together, these results indicate that ecDNA-chromosome tethering depends on active mitotic transcription of ecDNA-encoded genes like *PVT1* that are frequently co-amplified with *MYC*.

## DISCUSSION

Here, we show that ecDNA achieve high fidelity of nuclear inheritance by tethering to mitotic chromosomes. ecDNA exhibited high fidelity for nuclear segregation across a range of naturally occurring and artificially generated human cancer cell line models and across ecDNA harboring different oncogenes suggesting that accurate nuclear segregation may be an intrinsic feature of ecDNA. We further show that ecDNA resist blocks against mitotic transcription and are actively transcribed throughout mitosis. The precise mechanisms by which ecDNA resist global transcriptional suppression in mitosis are unknown, but are in line with their capacity for rampant transcription that derives from their more open chromatin conformation and participation in transcriptional hubs^11–14^. Inhibiting mitotic transcription by treatment with chemical agents interferes with ecDNA-chromosome tethering and results in cytosolic mis-segregation. Based on these observations, we propose that active transcription of ecDNA throughout mitosis links ecDNA to chromosomes via an RNA-mediated tether. We show that disrupting ecDNA-chromosome tethering results in ecDNA mis-segregation, chromosomal re-integration, and HSR formation. Thus, our work illustrates a new mechanism to explain how ecDNA-chromosome tethering preserves the high fidelity nuclear inheritance of ecDNA.

Although our data support an active model of ecDNA segregation, the process operates at an imperfect fidelity. Mis-segregating ecDNA were detected in micronuclei, where they undergo DNA damage and transcriptional silencing. Micronuclei are key platforms for chromosome rearrangement and cancer genome evolution, including catastrophic processes like the chromosome pulverizations seen in chromothripsis^3,5,6,8,35,53^. Aberrant resection and accumulation of single-stranded DNA in micronuclei may expose ecDNA to mutagenic processes like APOBEC3 deamination^54–56^. Ongoing ecDNA mis-segregation into micronuclei may, therefore, provide opportunities for continued ecDNA evolution. Indeed, ecDNA exhibit increased structural complexity at more advanced stages of disease^9^.

Our data indicate that BRD4 plays a critical role in protecting against cytosolic mis-segregation and chromosomal re-integration of ecDNA. These findings draw conceptual parallels to BRD4-dependent physical linkages that organize ecDNA into interphase hubs and anchor human papillomavirus genomes to mitotic chromosomes through an interaction with the viral E2 protein^13,57^. However, our data favor a model in which BRD4 safeguards ecDNA segregation through its canonical role in transcriptional co-activation. We can envision several models to explain how BRD4-dependent ecDNA transcription ensures nuclear inheritance. ecDNA-chromosome contacts bear resemblance to transcription hubs that form when distinct genomic loci are connected through physical interactions mediated in part by nascent RNA scaffolds^58–60^. Alternatively, persistent transcription of ecDNA in mitosis may be associated with the formation of RNA-DNA hybrids termed R-loops^61^. Indeed, ecDNA exhibit high levels of ssDNA accumulation, consistent with the presence of R-loops^15^. R-loops may therefore bring together different loci through an R-loop-anchored RNA bridge. Finally, by clustering and tethering to mitotic chromosomes, ecDNA bypasses normal mitotic processes, where the surfactant-like protein Ki-67 disperses chromosomes in early stages of mitosis to prepare for their individual segregation^62,63^. Instead, ecDNA resemble chromosomes during mitotic stages where co-condensation of Ki-67 with ribosomal RNAs at the perichromosomal layer endows Ki-67 with a chromosome attractant activity^64–67^. How Ki-67 affects ecDNA behavior during mitosis provides an avenue for further exploration into the mechanisms driving ecDNA nuclear inheritance. Similarly, the significance of the enrichment of ecDNA contact sites with the ends of mitotic chromosomes remains to be determined, but may be related to telomere transcription and production of telomeric repeat-containing RNA (TERRA)^68^.

HSR formation and gene amplification are thought to initiate when iterative processes, like breakage-fusion-bridge, capture sister chromatids to create a large palindrome^69^. Asymmetric breakage of this fused chromosome followed by repeated breakage-fusion-bridge cycling can then amplify genetic material over subsequent cell cycles^70^. In contrast, our data show that ecDNA to HSR conversion can occur on a timescale of hours that is inconsistent with separation by multiple cellular generations. These findings therefore support a model in which HSR formation is a rapid, all-at-once process driven through the simultaneous re-integration of multiple ecDNA copies into a single locus. Additionally, experimental induction of ecDNA to HSR conversion upon acute BRD4 inhibition and transient *PVT1* depletion provides an opportunity to study mechanisms underlying ecDNA reintegration and HSR formation. Future work will further investigate if ecDNA contacts with chromosome ends during mitosis drive preferential reintegration into these sites, as has been previously proposed^3^.

By demonstrating that ecDNA segregation is an active process that plays a critical role in ecDNA maintenance, our results point towards new avenues for treating ecDNA-containing cancers. Targeting ecDNA segregation may represent a promising strategy to prevent adaptive increases in ecDNA copy number that can occur following treatment with targeted therapies^15,22,24,71^. Strategies seeking to leverage the distinct properties of ecDNA segregation are likely to spare normal chromosome segregation and are thus likely to offer an improved therapeutic window.

### Limitations of the study

Our study primarily focuses on *MYC*-amplifying ecDNA endogenous to COLO320-DM and PC3 colorectal and prostate cancer cell lines. Although our data are consistent with the existence of mechanisms that actively promote the nuclear inheritance of non-*MYC*-amplifying ecDNA species, such as *EGFR*, *DHFR*, and *FGFR2*, it remains to be seen if similar transcription-based mechanisms or production of long non-coding RNAs are operative in these models. Furthermore, although *PVT1* depletion and transcription inhibition increase the rates of cytosolic mis-segregation, many ecDNA molecules continue to be segregated into the nucleus under these conditions. Therefore, we propose that additional mechanisms might ensure ecDNA nuclear segregation.

## ACKNOWLEDGEMENTS

We thank P. Mischel, H. Chang, V. Bafna, D. Cleveland, A. Sfeir, and A. Ventura for advice and reagents, the Epigenetics Research Innovation Lab and Integrated Genomics Operation at MSKCC for assistance with HiC analysis and shallow whole genome sequencing. Work in J.M.’s laboratory is supported by the NCI (R37CA261183; R01CA270102; P30CA008748), the Pershing Square Sohn Cancer Research Alliance, the Frank A. Howard Scholars Program, the Mary Kay Ash Foundation and the Experimental Therapeutics Centers at MSKCC. This material is based upon work supported by the National Science Foundation Graduate Research Fellowship under Grant No. 2234691 (A.N).

## AUTHOR CONTRIBUTIONS

Conceptualization: A.N and J.M.; Methodology: A.N. and J.M.; Investigation: A.N., R.N., Y.C., J.S., E.S.; Formal Analysis: Y.C. and R.K.; Writing - Original Draft: A.N. and J.M.; Writing - Review and Editing - all authors; Visualization: J.M., E.T., and A.N.; Funding Acquisition: A.N., R.K. and J.M; Supervision: R.K. and J.M.

## Supplementary Figures

**Figure S1.**
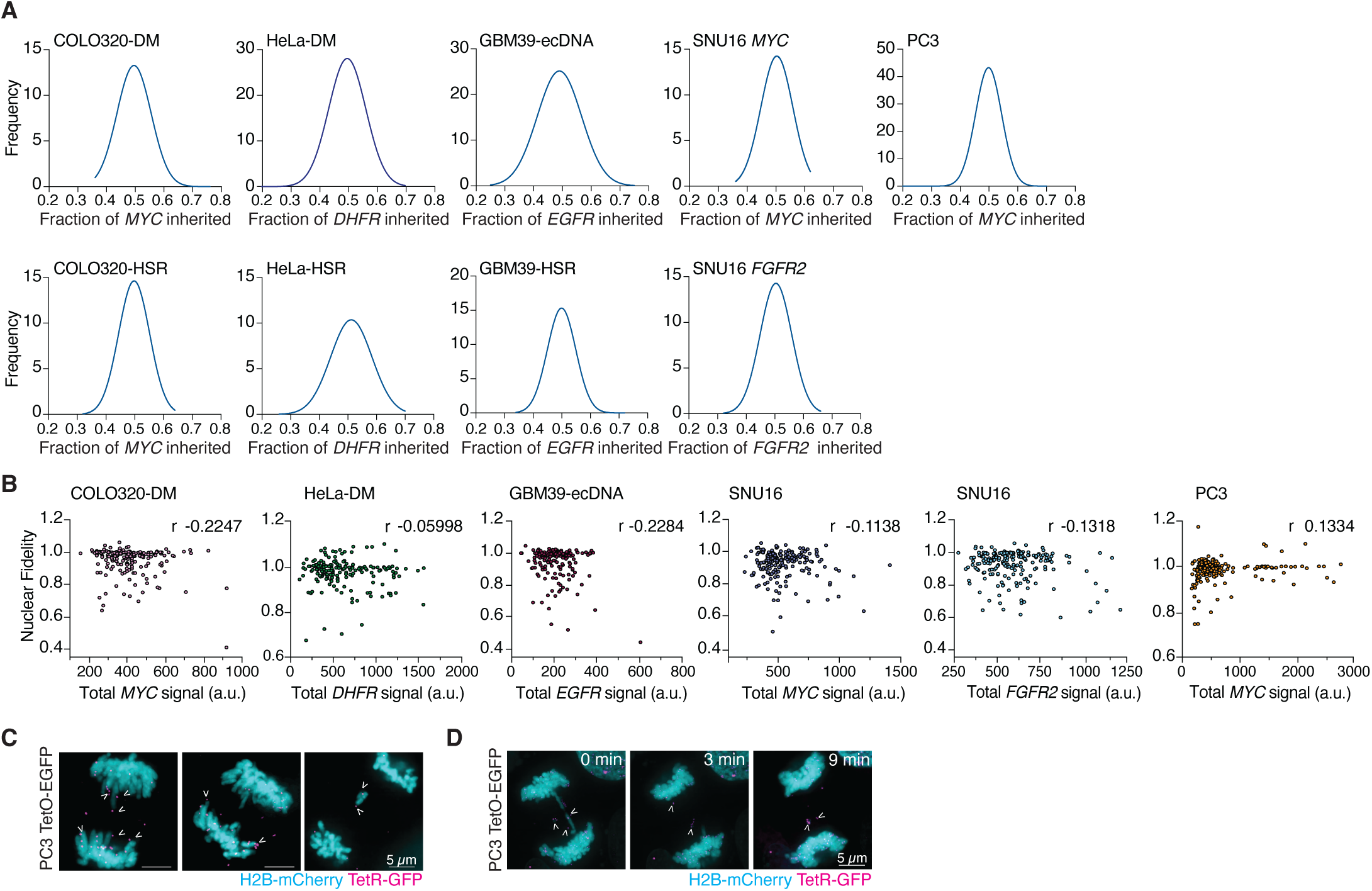
ecDNA asymmetric inheritance and nuclear segregation, related to Figure 1. (A) Frequency histogram of ecDNA inheritance in daughter cells following cell division for cancer cell panel in Figure 1A. Gaussian distributions are shown for each (*n =* 3 experiments, *n =* 100 pairs of daughter nuclei, >25 pairs analyzed per experiment). (B) Quantification of nuclear fidelity calculated in Figure 1B compared to ecDNA signal in each cell. Each point represents a single daughter cell. The Pearson Correlation Coefficient (*r)* is reported for each graph. (*n =* 3 experiments, *n =* 200 daughter nuclei) (C) Examples of live-cell imaging in PC3 TetO-EGFP cells expressing H2B-mCherry and TetR-GFP demonstrating ecDNA contacts with chromosome ends or sub-telomeric regions during mitosis. Arrows mark ecDNA making contacts with chromosome ends/subtelomeric regions. (D) Live-cell imaging of PC3 TetO-EGFP cells expressing H2B-mCherry and TetR-GFP. Insets mark time since anaphase onset (t = 0). Arrowheads mark ecDNA mis-segregating into the cytosol. All scale bars = 5µm.

**Figure S2.**
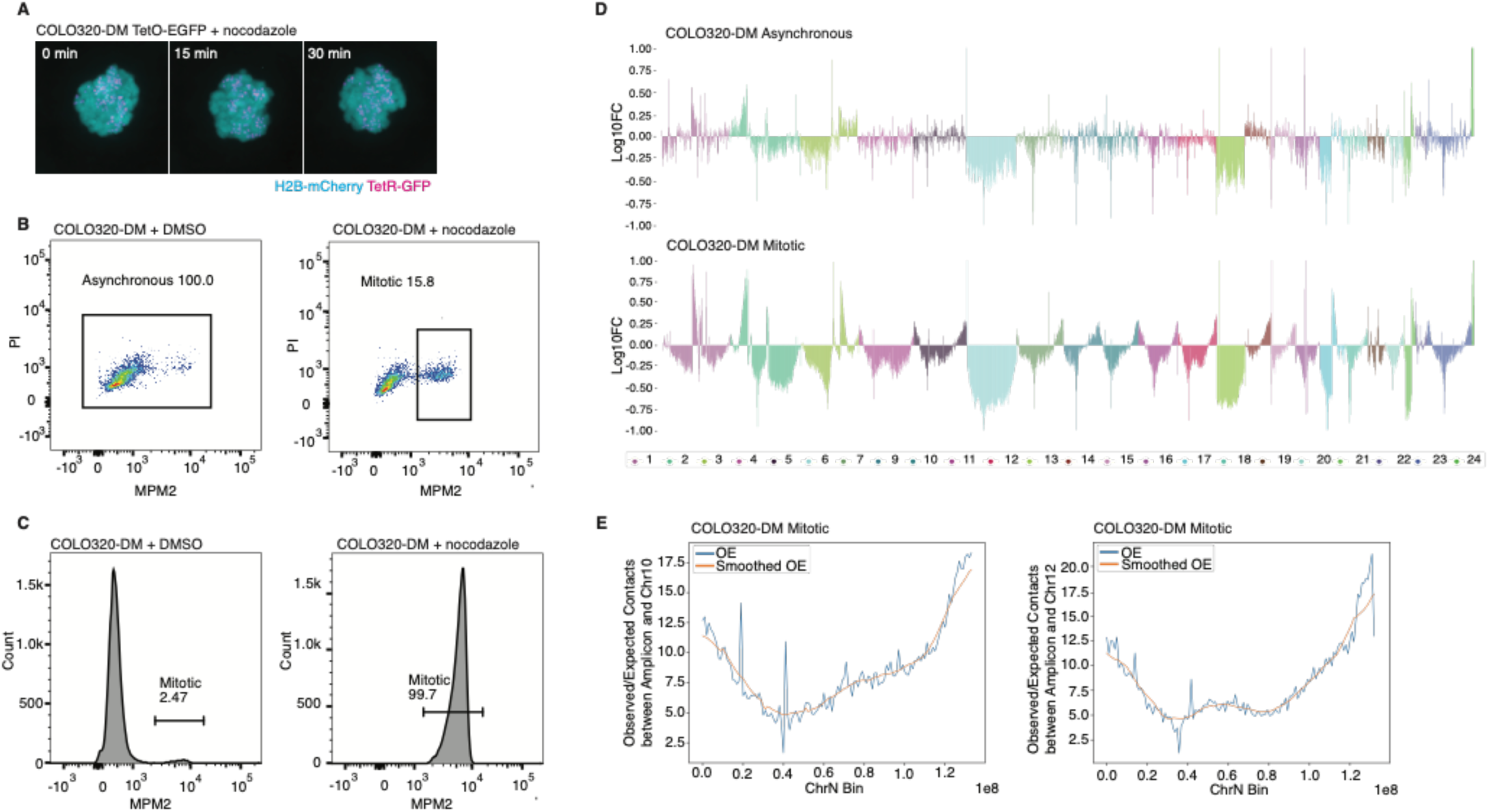
HiC sequencing of asynchronous versus mitotic COLO320-DMs, related to Figure 1. (A) Representative live-cell imaging of nocodazole-arrested COLO320-DM TetO-EGFP cells expressing H2B-mCherry and TetR-GFP. (B) Gating strategy for FACS sorting of asynchronous and mitotic samples for HiC sequencing. COLO320-DM cells were stained with an anti-MPM2 antibody and propidium iodide (PI). Single cells that were MPM2+PI+ were collected. (C) Post-sorting flow cytometry analysis of sorted COLO320-DM asynchronous versus mitotic cells. (D) Linear chromosome map demonstrating the log10 fold change in observed over valid expected (OE) contacts between the COLO320-DM amplicon and the genome in 1 Mb bins. Chromosome 8 has been removed. (E) Additional examples of linear chromosome maps obtained via HiC sequencing of COLO320-DM mitotic samples. Observed over valid expected (OE) contacts between the amplicon region and Chromosome 10 (left) and Chromosome 12 (right) are shown.

**Figure S3.**
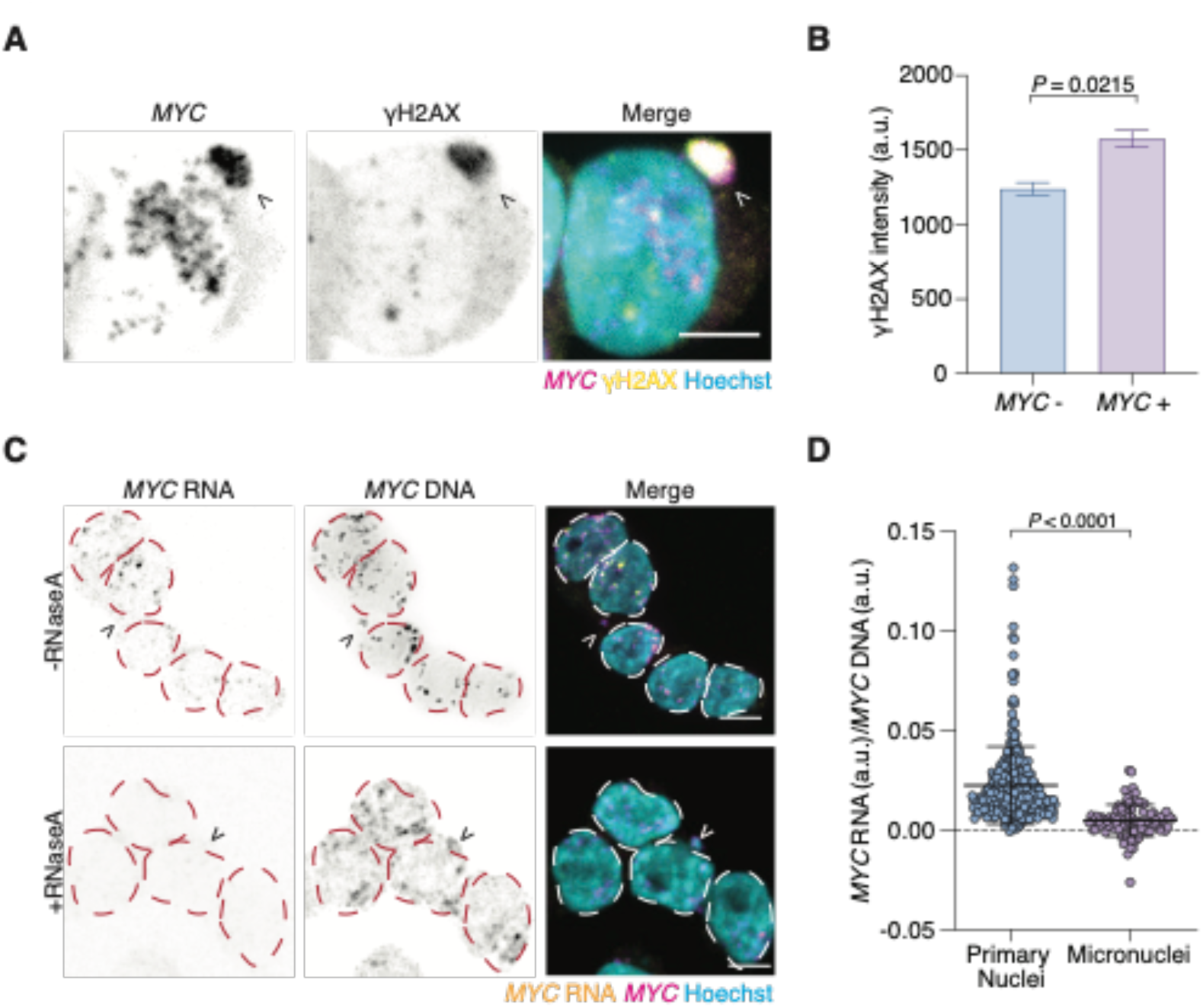
ecDNA mis-segregation into the cytosol, related to Figure 1. (A) Immunofluorescence-FISH staining of γH2AX and *MYC* ecDNA using a *MYC-*specific FISH probe in COLO320-DM. Arrowhead indicates ecDNA+ micronucleus. (B) Quantification of γH2AX signal intensity in *MYC*-versus *MYC*+ micronuclei in COLO320-DM cells. Mean±s.e.m. of *n* = 3 experiments are shown. (*n* = 164 micronuclei; >45 micronuclei per experiment). *P* value was calculated using an unpaired *t*-test. (C) Combined RNA/DNA FISH for *MYC* transcripts and *MYC* DNA in COLO320-DM cells. *MYC* transcripts are stained with an intron-specific RNA FISH probe. Borders of nuclei are highlighted, and arrows indicate *MYC*+ micronuclei. (D) Quantification of combined RNA/DNA FISH shown in (C). Mean±s.d. of *n* = 3 experiments (>100 primary nuclei analyzed per experiment) are shown. *P* values were calculated using unpaired *t*-test.

**Figure S4.**
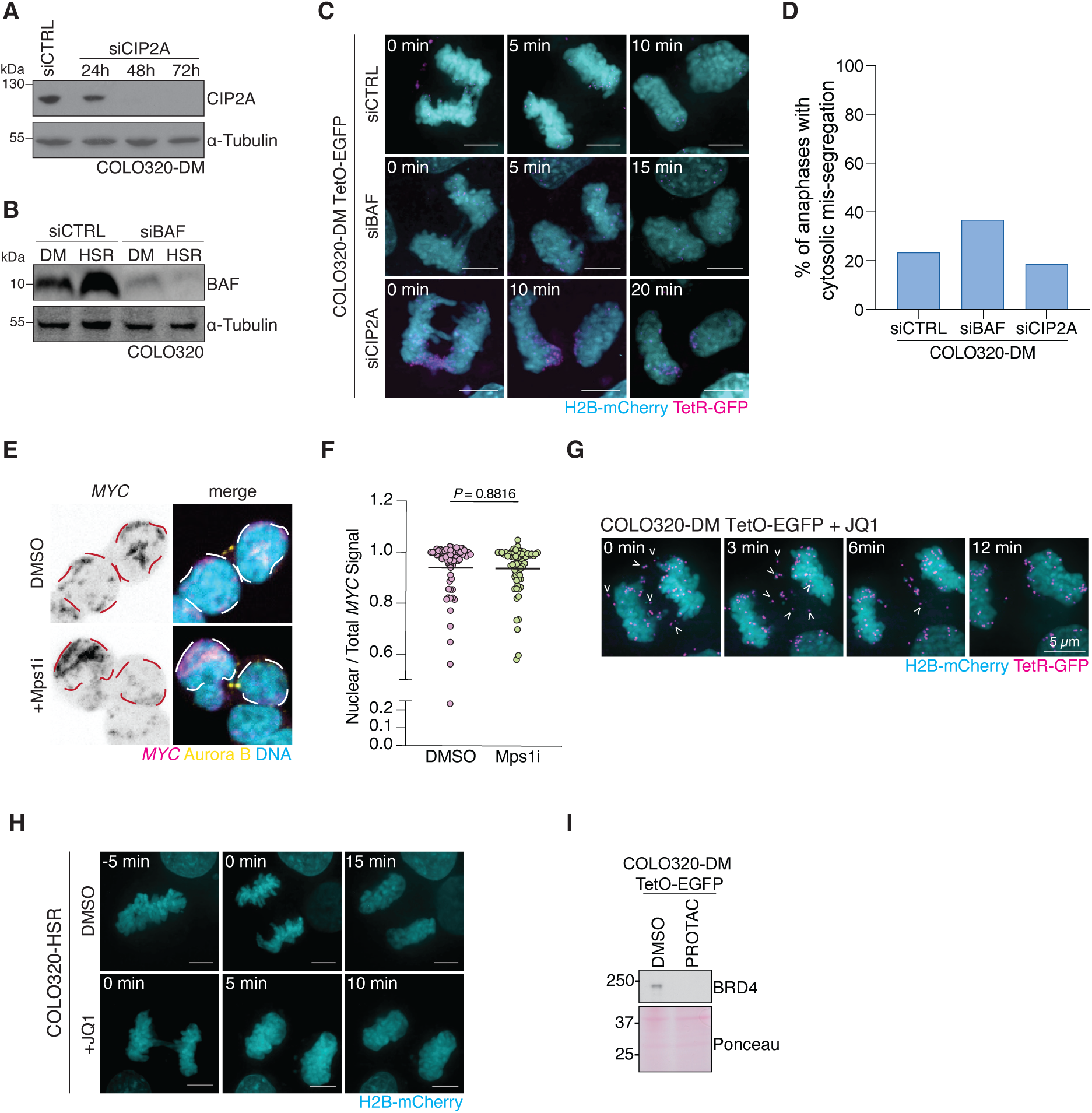
BRD4 inhibition drives ecDNA mis-segregation, related to Figure 2. (A) Immunoblotting for CIP2A and ɑ-tubulin in COLO320-DM cells transfected with siCTRL or siCIP2A for indicated time. (B) Immunoblotting for BAF and ɑ-tubulin in COLO320-DM and COLO320-HSR cells following transfection with siCTRL and siBAF for 72 hours. (C) Live-cell imaging of COLO320-DM TetO-GFP cells expressing H2B-mCherry and TetR-GFP following transfection with indicated siRNAs. Insets indicate time in minutes since anaphase onset (t *=* 0). (D) Quantification of ecDNA cytosolic mis-segregation from COLO320-DM TetO-EGFP cells transfected with siCTRL, siCIP2A, or siBAF, as shown in (C) (siCTRL *n* = 17, siBAF *n* = 19, siCIP2A *n* = 18). (E) Immunofluorescence-DNA FISH of COLO320-DM cells treated with DMSO or Mps1i for 24 hours. Recently divided cells were identified via anti-Aurora B immunofluorescence and ecDNA was identified using a *MYC* specific FISH probe. Nuclei borders are highlighted. (F) Quantification of ecDNA nuclear segregation fidelity in COLO320-DM cells treated with DMSO or MPS1i as shown in (E). Mean of *n* = 1 experiment is shown with a minimum of 25 pairs of recently divided cells analyzed. *P* values were calculated using unpaired t tests. (G) Live-cell imaging of COLO320-DM TetO-EGFP cells treated with 500 nM JQ1 for 6 hours. Insets indicate time in minutes since anaphase onset (t = 0) and arrows mark ecDNAs mis-segregating into the cytosol. (H) Live-cell imaging of COLO320-HSR cells expressing H2B-mCherry treated with DMSO or 500 nM JQ1 for 6 hours. Insets indicate time in minutes since anaphase onset (t = 0). (I) Immunoblotting of BRD4 in COLO320-DM TetO cells treated with DMSO or HaloPROTAC for 48 hours. Total protein is stained via Ponceau. Scale bars = 5µm.

**Figure S5.**
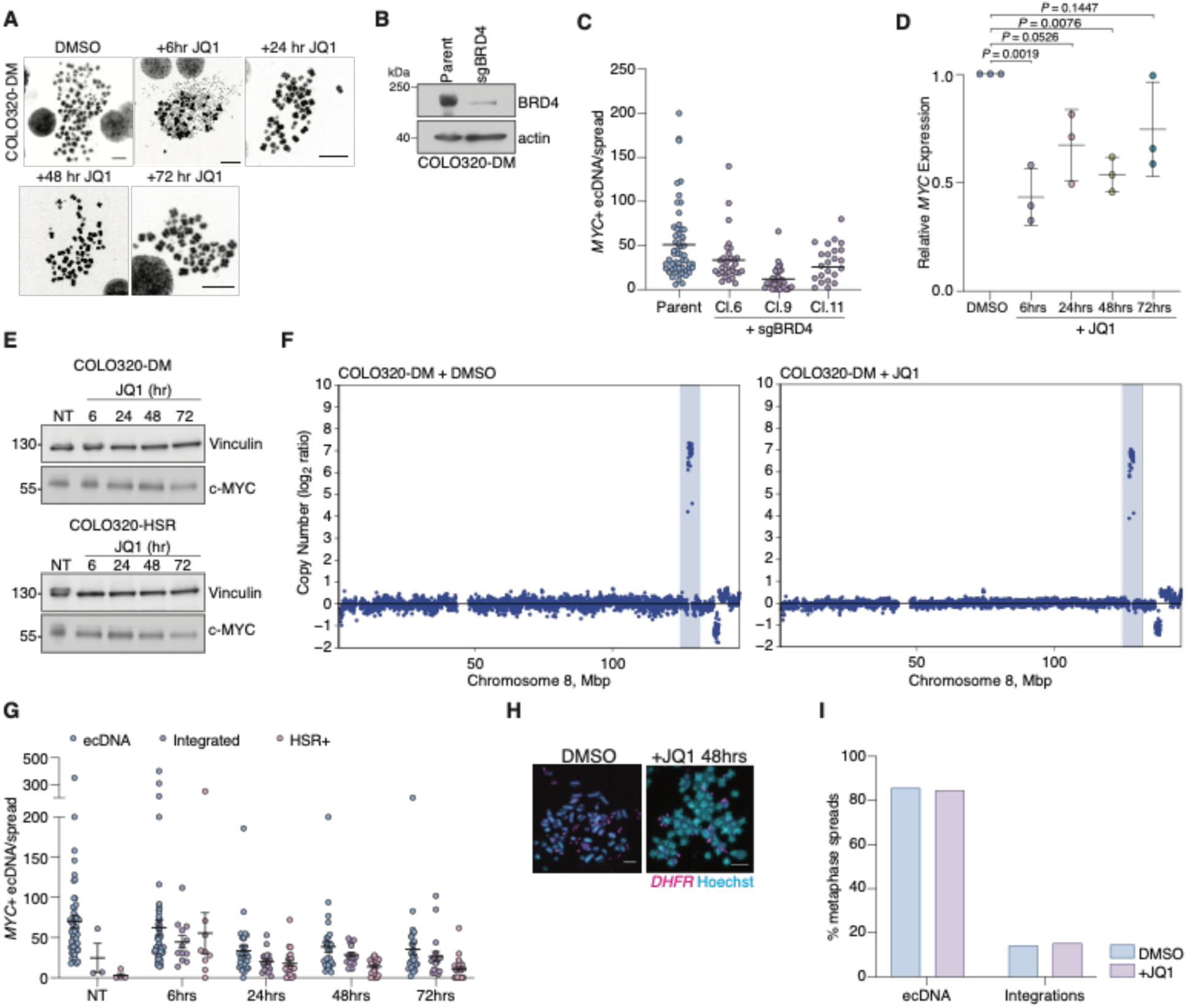
BRD4 inhibition drives HSR formation, related to Figure 2. (A) Representative metaphase spreads of COLO320-DM cells treated with 500 nM JQ1 for the indicated time. (B) Immunoblotting of BRD4 and actin in parental COLO320-DM cells or COLO320-DM cells transfected with Cas9 and sgRNAs specific for *BRD4*. (C) Metaphase spread analysis showing the number of ecDNA in COLO320-DM parental or COLO320-DM BRD4 hypomorph clones. Average ecDNA number shown of *n* = 1-2 experiments (>25 metaphase spreads per cell line analyzed). (D) Quantitative PCR analysis of *MYC* transcripts in COLO320-DM cells treated with 500 nM JQ1 for indicated time period. Mean±s.d. are indicated of *n* = 3 experiments. *P* values were calculated by one-way ANOVA with Tukey’s multiple comparisons test. (E) Immunoblotting of c-MYC and vinculin of COLO320-DM and COLO320-HSR cell lines treated with DMSO or 500 nM JQ1 for the indicated times. (F) Copy number analysis from shallow whole genome sequencing of COLO320-DM cells treated with DMSO or 500 nM for seven days. Highlighted region of Chromosome 8 indicates *MYC* sequence amplified on ecDNA in COLO320-DM cells. MBp = megabase pairs (G) Metaphase spread analysis showing number of ecDNA in COLO320-DM cells treated with 500 nM JQ1 for indicated times as shown in Figures 2H and 2I. Mean±s.e.m. are indicated. Spreads are grouped based on whether they demonstrated *MYC* only amplified on ecDNA (ecDNA), showed evidence of *MYC* integrations in the linear chromosomes (Integrations), or contained *MYC+* homogenously staining region(s) (HSR+). Mean±s.d. of *n* = 3 independent experiments (>25 metaphase spreads per condition were analyzed). (H) Representative metaphase *DHFR* FISH examples from HeLa-DM cells treated with DMSO or JQ1 for 48 hours. Average number of ecDNA from this experiment are quantified in Figure 2J. (I) Quantification of number of metaphase spreads from metaphase *DHFR* FISH shown in Figure 2J and Figure S4H demonstrating integration of *DHFR* into the linear chromosome. Unless otherwise noted, *P* values were calculated using unpaired *t*-tests. Scale bars = 5 µm.

**Figure S6.**
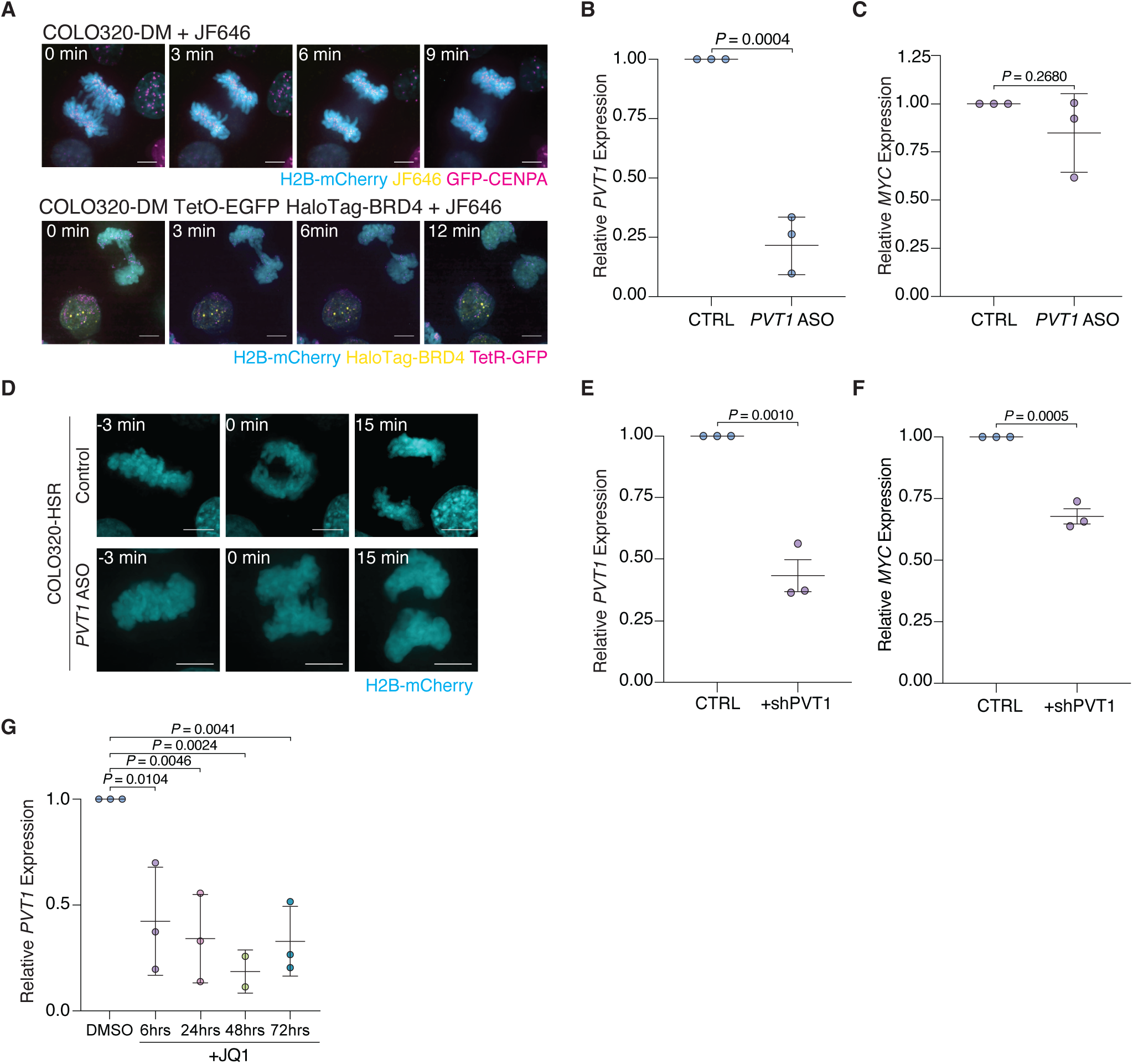
PVT1 knockdown by ASO, shRNA, and JQ1 treatment, related to Figure 3. (A) Live-cell imaging of COLO320-DM cells expressing H2B-mCherry and GFP-CENPA (top) or COLO320-DM TetO-GFP HaloTag-BRD4 cells (bottom) stained with JF646 ligand. Specific staining of HaloTag-BRD4 in COLO320-DM TetO-GFP confirmed by lack of foci in parent plus ligand control (top). Insets indicate time since anaphase onset (t = 0). (B) Quantitative PCR analysis of *PVT1* transcripts in COLO320-DM cells transfected with control or *PVT1* ASO for 48 hours. Mean±s.d. of *n* = 3 biological replicates are shown. (C) Quantitative PCR analysis of *MYC* transcripts in COLO320-DM cells transfected with control or *PVT1* ASO for 48 hours. Mean±s.d. of *n* = 3 biological replicates are shown. (D) Live-cell imaging of COLO320-HSR cells expressing H2B-mCherry transfected with control or *PVT1-*specific ASO for 48 hours before image acquisition. Insets indicate time in minutes since anaphase onset. (E) Quantitative PCR analysis of *PVT1* transcripts in COLO320-DM cells expressing shCTRL or shPVT1. Mean±s.d. of *n* = 3 biological replicates are shown. (F) Quantitative PCR analysis of *MYC* transcripts in COLO320-DM cells expressing shCTRL or shPVT1. (G) Quantitative PCR analysis of *PVT1* transcript levels in COLO320-DM cells treated with 500 nM JQ1 for indicated time period. Mean±s.d. of *n* = 3 biological replicates are shown. *P* values were calculated by one-way ANOVA with Tukey’s multiple comparisons test. Unless otherwise noted, *P* values were calculated using unpaired *t*-tests. Scale bars = 5 µm.

**Figure S7.**
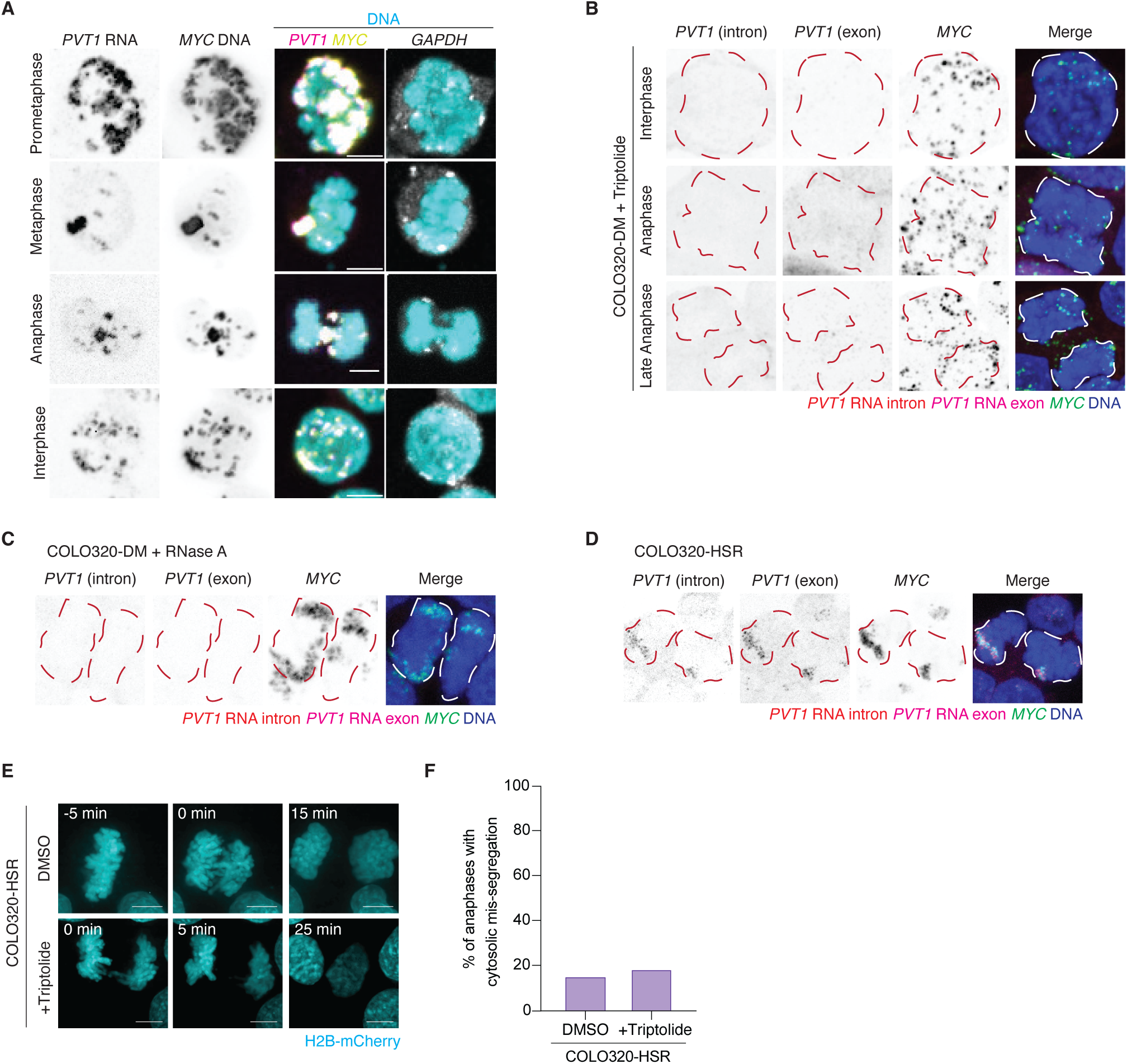
Transcription of ecDNA-encoded *PVT1* during mitosis, related to Figure 4. (A) Combined RNA/DNA FISH of COLO320-DM cells using PVT1 exon and GAPDH RNA FISH probes and *MYC* DNA FISH probe. Representative example images of interphase and mitotic COLO320-DM cells are shown. (B) Representative examples of RNA/DNA FISH in COLO320-DM cells treated with triptolide for 3.5 hours. *PVT1* transcripts were stained with an intron and exon probe, ecDNA were stained with a *MYC*-specific DNA FISH probe. Cells fixed in different stages of the cell cycle are depicted. Nuclei borders are indicated. (C) Representative example of RNA/DNA FISH in COLO320-DM cells treated with RNase A for 2 hours. *PVT1* transcripts were stained with an intron and exon probe, ecDNA were stained with a *MYC* specific DNA FISH probe. Nuclei borders are highlighted. (D) Representative example of RNA/DNA FISH in COLO320-HSR cells. *PVT1* transcripts were detected with intronic and exon probes, ecDNA were stained with a *MYC*-specific DNA FISH probe. Nuclei borders are indicated. (E) Live-cell imaging of COLO320-HSR cells expressing H2B-mCherry treated with DMSO or triptolide for 2 hours before starting image acquisition. Insets indicate time since anaphase onset (t = 0). (F) Quantification of anaphases from live-cell imaging shown in (E) of COLO320-HSR cells treated with DMSO or triptolide (DMSO *n* = 20; Triptolide *n* = 22). Scale bars = 5 µm.

**Video S1.** Live-cell imaging of untreated COLO320-DM TetO-GFP cells expressing H2B-mCherry and TetR-GFP.

**Video S2.** Live-cell imaging of untreated PC3 TetO-EGFP cells expressing H2B-mCherry and TetR-GFP.

**Video S3.** Live-cell imaging of COLO320-DM TetO-EGFP cells expressing H2B-mCherry and TetR-GFP treated with DMSO.

**Video S4.** Live-cell imaging of COLO320-DM TetO-EGFP cells expressing H2B-mCherry and TetR-GFP treated with JQ1.

**Video S5.** Live-cell imaging of COLO320-DM TetO-EGFP cells expressing H2B-mCherry and TetR-GFP treated with triptolide.

## STAR Methods

### KEY RESOURCES TABLE

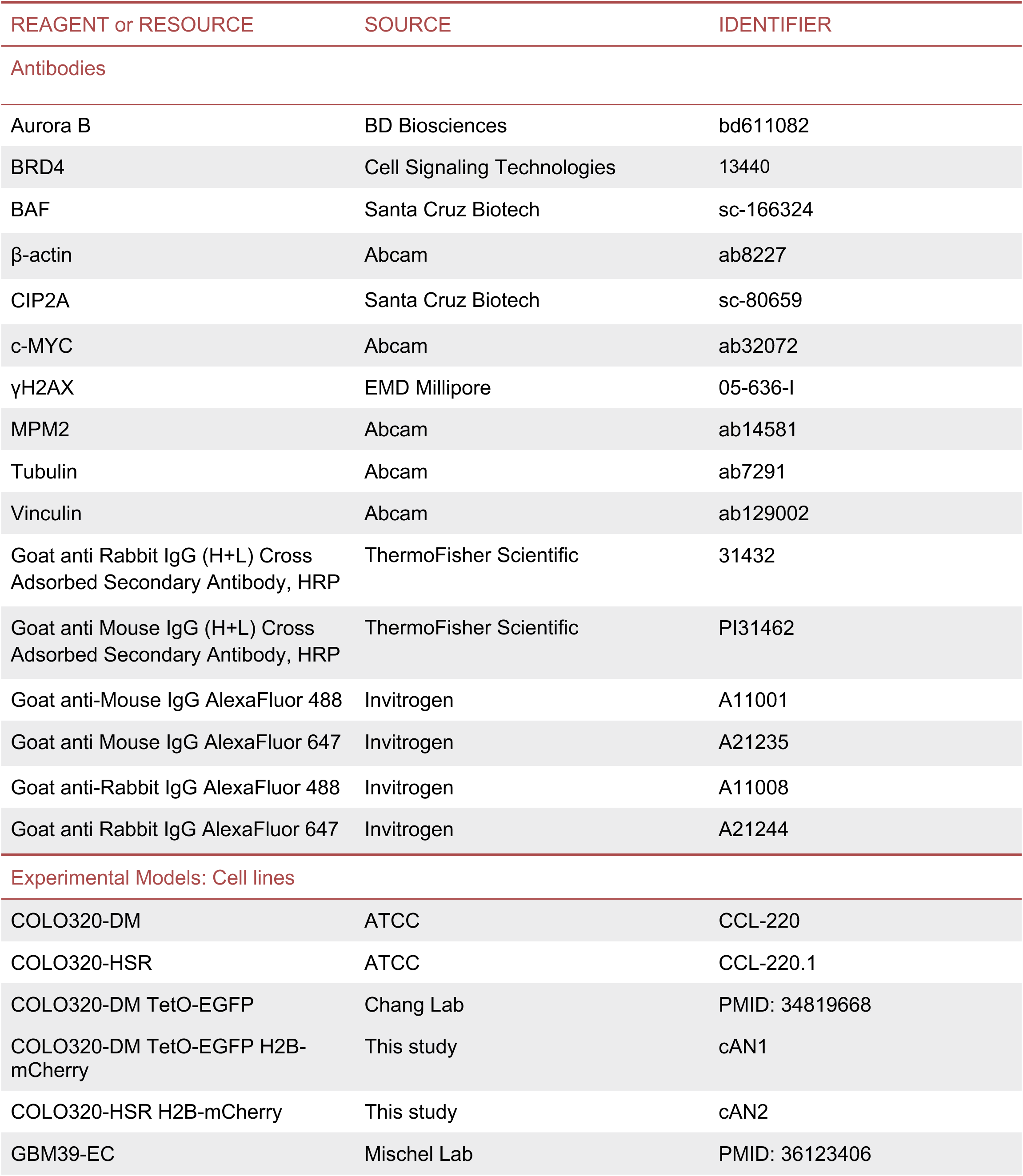
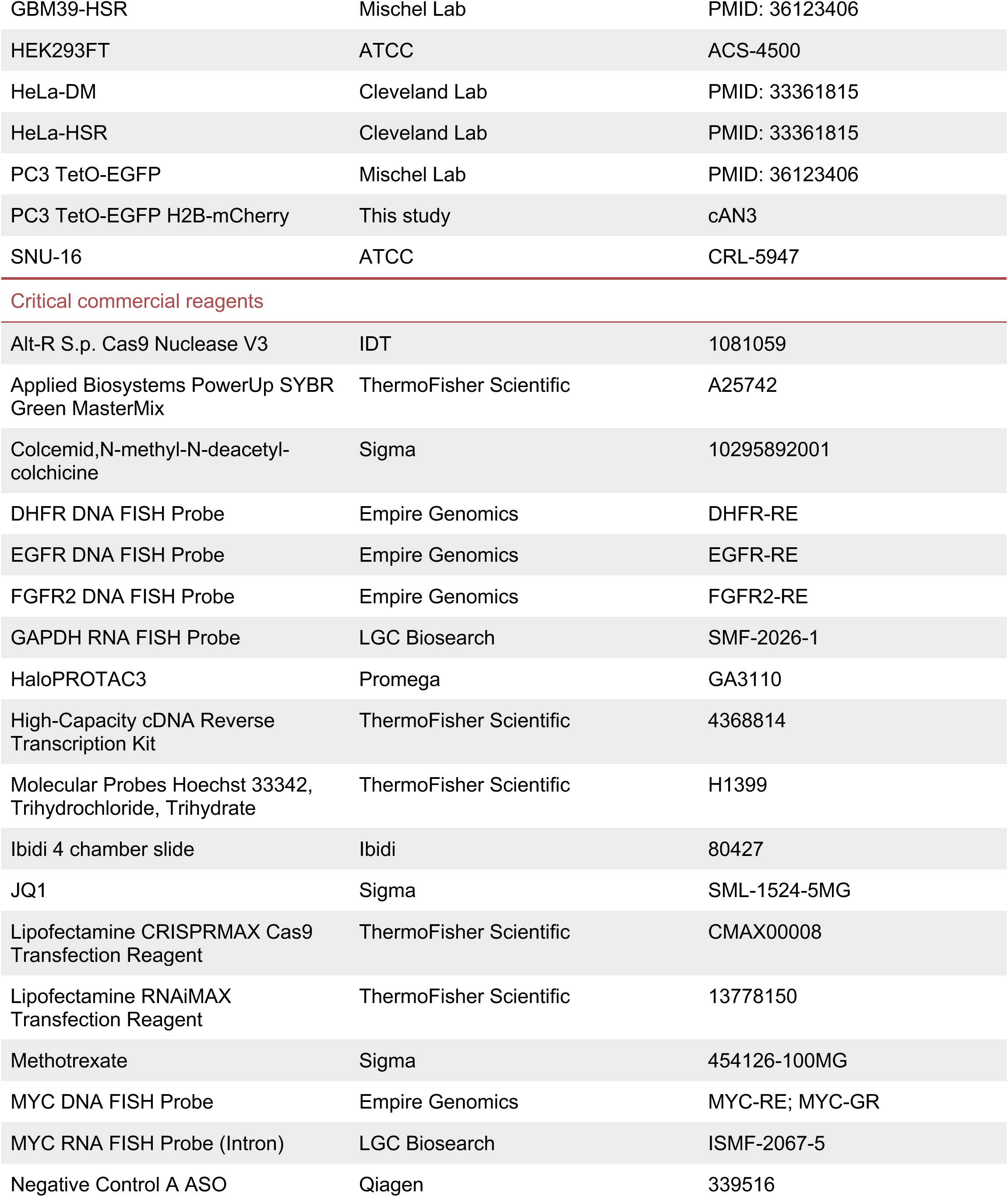
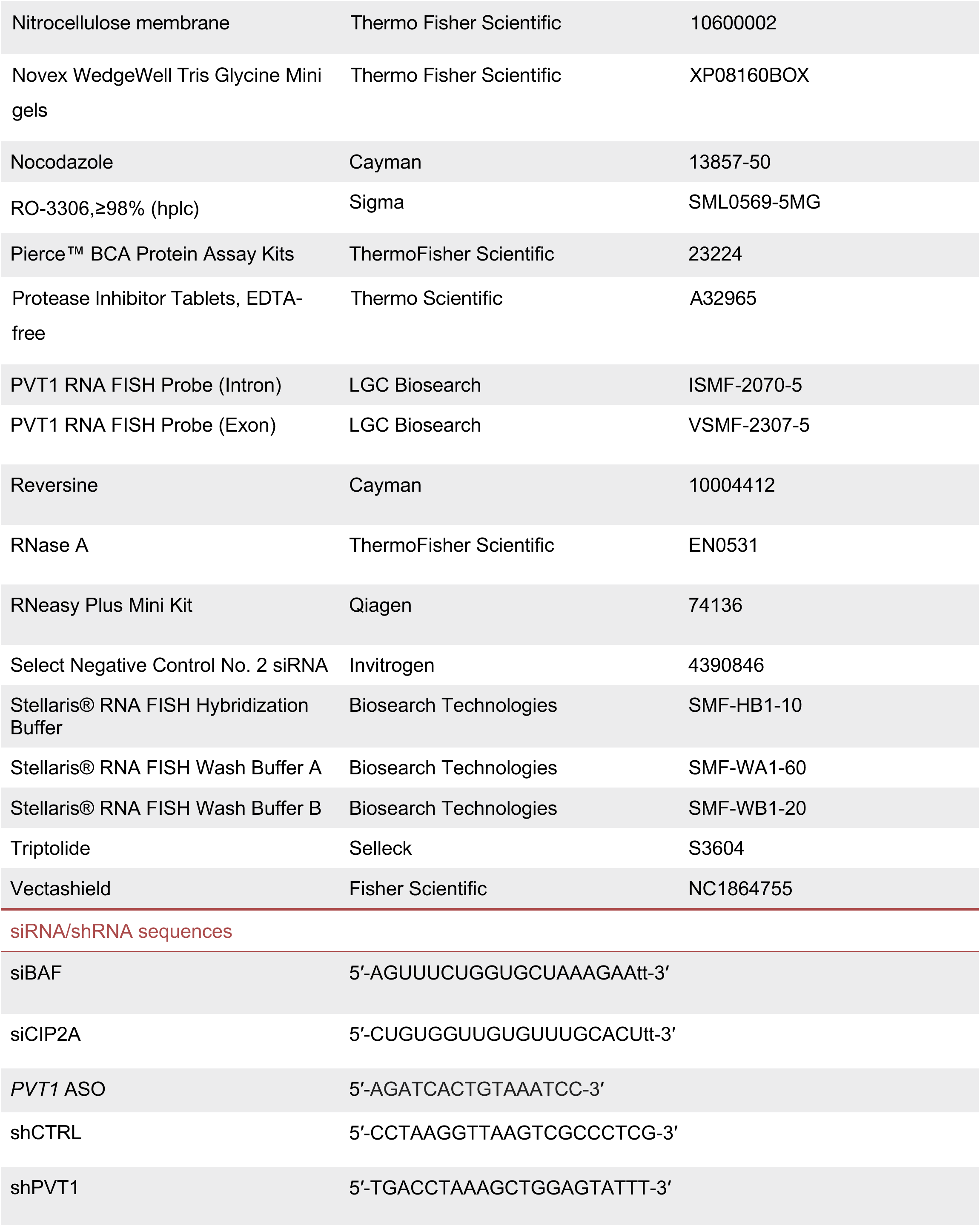
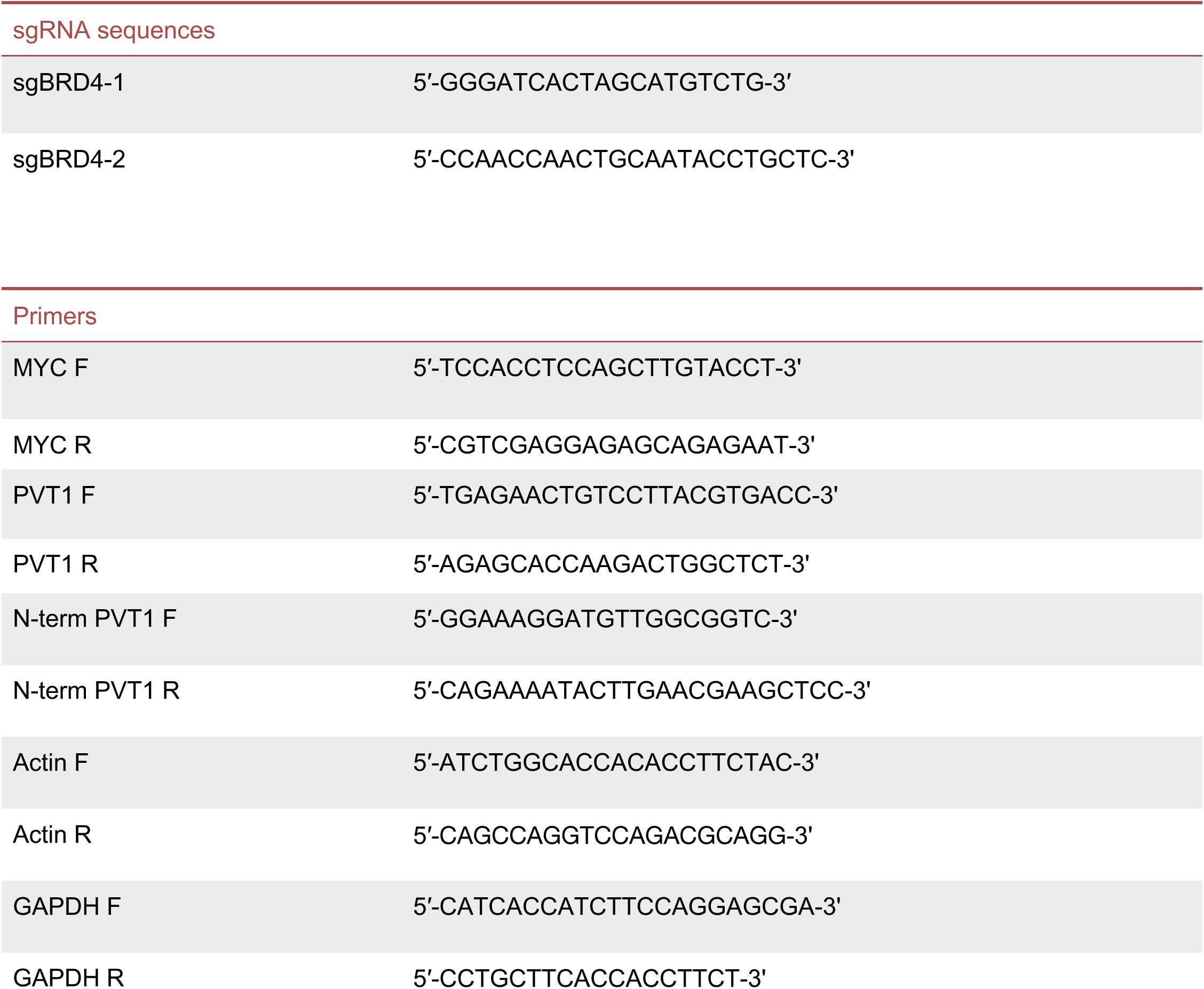

### RESOURCE AVAILABILITY

#### Lead contact

Further information and requests for resources and reagents should be directed to and will be fulfilled by the Lead Contact, John Maciejowski.

#### Materials availability

Cell lines generated in this study and listed in the Key Resource Table are available from Dr. John Maciejowski.

#### Data and code availability

The datasets and original source data generated during this study are available

- All data reported in this paper will be shared by the lead contact upon request.
- The paper does not report original code.

### EXPERIMENTAL MODEL DETAILS

#### Cell culture

COLO320-DM (ATCC; catalog no. CCL-220), COLO320-HSR (ATCC; catalog no. CCL-220.1), SNU16 (ATCC; catalog no. CRL-5974) cell lines were purchased from the American Type Culture Collection (ATCC). COLO320-DM COLO320-DM, COLO320-HSR, and SNU-16 cell lines were cultured in RPMI supplemented with 10 % FBS.

TetO-EGFP COLO320-DM cell line was a gift from the Howard Chang lab. TetO-EGFP PC3, GBM39-EC, and GBM39-HSR cell lines were a gift from the Paul Mischel lab. TetO-EGFP PC3 cells were cultured in DMEM supplemented with 10 % FBS and 1 % penicillin-streptomycin. GBM39-EC and GBM39-HSR cell lines were cultured in 1:1 mixture of F12:DMEM media supplemented with 1 % GlutaMax (Gibco), 10 % B-27 serum-free supplement (Gibco), 20 ng/mL human EGF (Sigma), 20 ng/mL human FGF (Sigma), and 1 μg/mL heparin (Sigma). Media was supplemented with 1% penicillin-streptomycin.

HeLa-DM and HeLa-HSR cell lines were a gift from the Don Cleveland lab. HeLa-DM and HeLa-HSR correspond to MTX-resistant clone PD29425G and MTX-resistant clone PD29427P, as described previously^1^. HeLa-DM and HeLa-HSR cell lines were cultured in DMEM supplemented with 10% dialyzed FBS and 1% penicillin-streptomycin. Cells were maintained in 80 nM methotrexate dissolved in DMSO (Sigma). Unless otherwise noted, all media and supplements were supplied by the MSKCC media core facility. All cell lines used in this study tested negative for mycoplasma.

Polyclonal COLO320-DM *BRD4* knockout cells were generated by transfecting Cas9 protein, BRD4-sgRNA1, and BRD4-sgRNA2 using Lipofectamine CRISPRMAX Cas9 Transfection Reagent (Invitrogen) per manufacturer’s instructions, then isolating and expanding hypomorphs from single cell colonies.

#### Pharmacological treatments

For HiC-sequencing mitotic cell preparation described in Figure 1 and Figure S2, COLO320-DM, COLO320-HSR, or COLO320-DM TetO-EGFP cells were incubated with 100 ng/mL nocodazole in DMSO (Cayman) for four hours prior to fixation. For live-cell imaging in Figure S2A, COLO320-DM TetO-EGFP cells were treated with 100 ng/mL nocodazole for four hours before the start of imaging.

The Mps1 inhibitor, reversine (Cayman), was resuspended in DMSO to a concentration of 0.5 mg/mL. COLO320-DM cells were treated with reversine at a final concentration of 100 ng/mL for 24 hours prior to fixation (Figure S4E-F).

COLO320-DM and HeLa-DM cell lines, cells were incubated with JQ1 (Sigma) dissolved in DMSO for 6-72 hours at a final concentration of 500 nM (Figures 2A-B, Figures 2G-L, Figure S5A, Figure S5D-I, and Figure S6G). For live-cell imaging experiments in Figures 2C-F, Figure S4G, and Figure S4I, COLO320-DM TetO-EGFP, COLO320-HSR, and PC3-TetO-EGFP were incubated with 500 nM JQ1 in DMSO for 6 hours prior to imaging. In Figures 2C-D and Figure S4H, HaloTag-BRD4 degradation in COLO320-DM TetO-GFP was conducted via incubation of cells with 500 nM HaloPROTAC (Promega) for 48 hours prior to live-cell imaging. For co-treatment of JQ1 and RO-3306 in Figure 2K and 2L, 500 nM JQ1, and 9 µM RO-3306 (Sigma) were added simultaneously to COLO320-DM cells for 48 hours. To visualize HaloTag-BRD4 in COLO320-DM TetO-EGFP cells as shown in Figure S6A, cells were stained with Janelia Fluor HaloTag fluorescent ligand, JF646 (Janelia) at a final concentration of 200 nM. To minimize background, cells were incubated with JF646 the night before imaging for 15 minutes. Cells were then washed five times with PBS and incubated in fresh media overnight.

Triptolide (Selleck) was used to inhibit global transcription in COLO320-DM, COLO320-HSR, COLO320-DM TetO-EGFP, and PC3 TetO-EGFP cells. For RNA/DNA FISH imaging in Figures 4A-C and Figure S7B-D, COLO320-DM, and COLO320-HSR cells were incubated with triptolide at a final concentration of 5.5 µM for 3.5 hours prior to fixation. For Figures 4D-H and Figure S7E and S7F, COLO320-DM TetO-EGFP cells were treated with triptolide for 2 hours before live-cell imaging. PC3 TetO-EGFP cells were treated with triptolide for 4 hours prior to the start of imaging.

#### siRNA and ASO transfections

Custom siRNAs targeting BAF and CIP2A were ordered from Dharmacon ^2,3^. Silencer Select Negative Control No. 2 was used as siCTRL (See Key Resources Table). All siRNAs were transfected at a final concentration of 10 nM using Lipofectamine RNAiMAX Transfection Reagents (ThermoFisher Scientific), except for siBAF, which was transfected at a final concentration of 100 nM. COLO320-DM cells were transfected with siCTRL and siCIP2A for 48 hours before immunoblotting (Figure S3A) and live-cell imaging (Figure S4C and S4D). Cells were transfected with siCTRL and siBAF for 72 hours prior to immunoblotting (Figure S4B) and live-cell imaging (Figure S4C and S4D).

For antisense oligonucleotide (ASO) knockdown of *PVT1*, custom Antisense LNA GapmeRs were ordered from Qiagen along with the Antisense LNA GapmeR Negative Control A (See Key Resources Table). ASOs were resuspended in nuclease-free TE buffer for a final concentration of 50 µM. ASOs were transfected into COLO320-DM, COLO320-DM TetO-EGFP, and PC3 TetO-EGFP cells at a final concentration of 50 nM using Lipofectamine RNAiMAX Transfection Reagents (ThermoFisher Scientific) for 48 hours prior to harvest for analysis (Figure S6B and S6C; Figure S6E-G) or live-cell imaging (Figures 3A-D).

#### Lentivirus production and infection

3 × 10^6^ HEK293-FT cells were seeded into 10-cm dishes. Constructs were transfected into cells with psPAX2 (Addgene #12260) and pMD2.G (Addgene #12259) using calcium phosphate precipitation. Lentivirus-containing supernatants were collected and filtered. At the time of infection, the lentiviral supernatant was supplemented with 4 μg/mL polybrene. Successfully transduced cells were selected through fluorescence-activated cell sorting by FACSAria (BD Biosciences) at the MSKCC Flow Cytometry Core Facility.

For lentiviral overexpression of shCTRL and shPVT1, custom shRNA-containing lentiviral constructs were ordered from VectorBuilder with a BFP selection marker. Lentivirus was generated as described above, and cells were sorted for BFP expression by FACSAria before being harvested for gene expression analysis (Figure S6E and S6F) and metaphase spreads (Figures 3E-G).

### Metaphase chromosome spreads

Cells were arrested in metaphase via an overnight treatment with 9 µM RO-3306 (Sigma) followed by a washout into 0.1 µg/mL colcemid (Sigma) for 4 hours. Cells were washed with trypsin and collected into a single cell suspension, followed by a 30 minute incubation at 37 °C in pre-warmed 75 mM KCl. Cells were then fixed with ice cold Carnoy’s fixative (3:1 methanol: glacial acetic acid) and stored at 4 °C long term. Chromosome spreads were dropped onto humidified glass slides and washed three times with ice cold fixative. Cells were then stained with 5 μg/mL Hoechst diluted 1:2000 (ThermoFisher Scientific) for 12 minutes at room temperature and mounted with Vectashield (Fisher Scientific), or unstained slides were stored at −20 °C for future use.

### DNA FISH

Chromosome spreads were aged overnight at −20 °C. The following day, slides were equilibrated in PBS at room temperature and washed once in 2X SSC buffer. Slides were dehydrated in an ascending ethanol series (70 %, 85 % and 100 %) for approximately 2 minutes each. After being allowed to dry completely, Empire Genomics FISH probes were diluted 1:10 in hybridization buffer and applied to a microscope slide with a coverslip. Coverslips were sealed with rubber cement, and slides were denatured at 80 °C for 8 minutes. FISH probes were hybridized overnight at 37 °C in a dark, humidified chamber. Samples were then washed in 0.4X SSC at 65 °C for 2 minutes and 2X SSC 0.1 % Tween-20 at room temperature for 30 sec. DNA was stained with Hoechst (5 μg/mL) diluted 1:2000 for 12 minutes at room temperature before being washed three times with PBS. Samples were briefly rinsed with dH_2_O. Coverslips were mounted with Vectashield and sealed with nail polish.

#### Immunofluorescence-DNA FISH

COLO320-DM, COLO320-HSR, PC3, HeLa-DM, and HeLa-HSR cells were grown on poly-L-lysine coated coverslips. SNU16 cells were plated on fibronectin coated coverslips (Neuvitro). GBM39-EC and GBM39-HSR cells were grown on laminin coated coverslips. 3.0 × 10^5^ cells/mL cells were plated on coverslips about 36 hours before fixation. Cells were washed once in PBS and fixed with 2 % PFA for 12 minutes at room temperature. Cells were washed once in PBS and stored in PBS with 0.02 % sodium azide at 4 °C.

Cells were permeabilized with 0.5 % Triton X-100 in PBS for 10 minutes at room temperature, then washed with PBS once. Samples were blocked in blocking buffer (1 mg/mL BSA, 3 % goat serum, 0.1 % Triton X-100, 1 mM EDTA in PBS) for 30 minutes at room temperature. Samples were incubated in primary antibody for 1-2 hours diluted 1:100 in blocking buffer at room temperature. Following primary antibody incubation, coverslips were washed three times in 0.05 % Triton X-100 in PBS. Cells were incubated in secondary antibody diluted 1:1000 in blocking buffer for 1 hour at room temperature in the dark. All subsequent steps were completed in the dark. After 1 hour, samples were washed three times with 0.05 % Triton X-100 in PBS. Cells were washed once with PBS and refixed with cold 2 % PFA for 20 minutes at room temperature. Coverslips were washed once in PBS and once in 2X SSC. FISH was carried out as described above with the following differences: Empire Genomics FISH probes were diluted 1:5 in hybridization buffer, and denaturation was conducted at 80 °C for 15-20 minutes.

#### RNA/DNA co-FISH

5.0 × 10^5^ cells/mL cells were plated on poly-L-lysine coated coverslips for 36-48 hours before fixation. Cells were fixed in 2 % PFA at room temperature, followed by a PBS wash and storage in PBS + 0.02 % sodium azide at 4 °C.

Cells were washed once in PBS at room temperature, followed by permeabilization in 0.5 % Triton X-100 in PBS for 10 minutes at room temperature. Cells were washed once with PBS and once in 2X SSC for 2 minutes at room temperature. Coverslips were then dehydrated in an ascending ethanol series (70 %, 85 % and 100 %) for 2 minutes each. Once completely dry, Empire Genomics DNA FISH probe was resuspended 1:5 in hybridization buffer and added to the coverslip and slide. Coverslips were sealed with rubber cement and denatured at 80 °C for 12-15 minutes. FISH probes were hybridized overnight at 37 °C in a humidified chamber in the dark. The following day, DNA FISH probes were washed off coverslips with 0.4X SSC at 65 °C and 2X SSC 0.1 % Tween-20 at room temperature for 2 minutes each. Coverslips were rinsed briefly with dH_2_O.

Coverslips were washed once in Stellaris RNA FISH Wash Buffer A at 37 °C (prepared according to manufacturer’s recommendation). Stellaris RNA FISH probes were resuspended in TE buffer to a concentration of 12.5 µM. For RNA FISH staining, RNA FISH probes were resuspended in Stellaris RNA FISH hybridization buffer (prepared according to manufacturer’s recommendation) 1:1000 for a final probe concentration of 125 nM and thoroughly mixed. RNA FISH probes were applied to coverslips and hybridized overnight in a dark, humidified chamber at 37 °C. For samples treated with RNaseA, recombinant RNAse A was diluted 1:100 in Stellaris RNA FISH Wash Buffer A. Coverslips were incubated with RNAse A in a dark, humidified chamber at 37 °C for 2-3 hours prior to hybridization with RNA FISH probe. Following incubation, coverslips were washed once with Stellaris Wash Buffer A before RNA FISH probes were applied.

After hybridization, RNA FISH probes were removed, and coverslips were washed once with Wash Buffer A for 30 minutes at 37 °C in the dark. Cells were counterstained with Hoechst (5 μg/mL) for 12 minutes at room temperature diluted in Wash Buffer A. Cells were washed once in Stellaris RNA FISH Wash Buffer B for 5 minutes at room temperature, rinsed in dH_2_O and allowed to dry. Coverslips were mounted with Vectashield and sealed with nail polish.

#### Immunoblotting

For Western Blots in Figure S4A, S4I, S5B, and S5E, 1 × 10^6^ cells were harvested by trypsinization and lysed in RIPA buffer (150 mM NaCl, 50 mM Tris-HCl, pH 8, 1 % NP-40, 0.5 % sodium deoxycholate, 0.1 % SDS, Pierce Protease Inhibitor Tablet, EDTA free) with 0.5mM PMSF. Cells were incubated in RIPA for 20 minutes on ice, and lysates were sonicated using a Bioruptor 3000 (Diagenode) for 15 cycles of 30 seconds ON/30 seconds OFF at 4 °C. Lysates were cleared by centrifugation at 20,000 × g for 20 minutes at 4 °C, and the protein concentration was quantified using the Pierce BCA Protein Assay kit (ThermoFisher Scientific). 20-30 µg of protein was loaded into Tris-Glycine gels (ThermoFisher Scientific). Protein transfer was performed by wet transfer with 1X Towbin Buffer (25 mM Tris, 192 mM glycine, 0.01 % SDS, 20 % methanol) and nitrocellulose membranes. Membranes were blocked in 5 % milk in 1X TBS-T (19 mM Tris, 137 mM NaCl, 2.7 mM KCl, and 0.1 % Tween-20) for 1 hour at room temperature and incubated with primary antibody diluted 1:1000 in blocking buffer (See Key Resources Table) overnight at 4 °C. Membranes were washed four times in 1X TBS-T followed by incubation with horseradish-peroxidase (HRP)-conjugated secondary antibodies (See Key Resources Table) diluted 1:10,000 in blocking buffer for 1 hour at room temperature. After four washes in 1X TBS-T, membranes were rinsed in TBS and visualized using enhanced chemiluminescence (ThermoFisher Scientific).

For BAF immunoblotting shown in Figure S3B, cells were harvested by trypsinization and lysed in RIPA buffer (150 mM NaCl, 50 mM Tris-HCl, pH 8, 1 % NP-40, 0.5 % sodium deoxycholate, 0.1 % SDS, Pierce Protease Inhibitor Tablet, EDTA free) with 0.5 mM PMSF. DNA was sheared 10 times with a 28 1/2 gauge insulin needle. Lysate was quantified by Pierce™ BCA Protein Assay Kits (ThermoFisher Scientific). 50 µg protein was loaded on 12 % Bis-Tris protein gels (Invitrogen, NP0342BOX). Gels were transferred to 0.2 μm PVDF membrane (Thermo Fisher, 88520) at 50V for 1 hour. Membranes were blocked in 1X AdvanBlock-Chemi blocking solution (Advansta, R-03023-D20) for 1 hour at room temperature and incubated with primary antibody diluted in blocking buffer overnight at 4 °C, washed 4 times in TBS-T, and incubated for 1 hour at room temperature with horseradish-peroxidase-conjugated secondary antibody. anti-BAF antibodies were diluted 1:500 in blocking buffer, and anti-β-actin primary antibodies were diluted 1:2000. After four washes in TBS-T, membranes were detected with Pierce ECL Western Blotting Substrate (Thermo Scientific) supplemented with 10 % Lumigen ECL Ultra (Lumigen) and imaging was performed using enhanced chemiluminescence (Amersham Image Quant 800 and BioRad ChemiDoc MP).

#### Gene expression analysis

For mRNA quantification in Figure S5D, S6B-C, and S6E-G, total RNA was isolated from 1 × 10^6^ cells using the RNeasy PLUS Mini RNA Isolation Kit (Qiagen) according to manufacturer’s instructions. RNA was quantified and converted to cDNA using the High Capacity cDNA Reverse Transcription Kit (ThermoFisher Scientific). Quantitative PCR was then performed using the Applied Biosystems PowerUp SYBR Green Master Mix (ThermoFisher Scientific), and detected on a QuantStudio6 (Applied Biosystems) cycler with gene-specific primers (See Key Resources Table). Relative transcript levels were determined by normalizing to both *GAPDH* and *Actin* expression levels using the ΔΔCt method on the Applied Biosystems Quantitative PCR Design & Analysis software.

#### Confocal microscopy

Images in Figure 1A, Figure 1E, Figure 2A, Figure 2H, Figure 3E, Figure 4A, Figure S3A, Figure S3E, Figure S4E, Figure S5A, Figure S5H, and Figure S7A-D were acquired on a Nikon Eclipse Ti2-E equipped with a CSU-W1 spinning disk with Borealis microadapter, Perfect Focus 4, motorized turret and encoded stage, polycarbonate thermal box, 5 line laser launch [405 (100 mw), 445 (45 mw), 488 (100 mw), 561 (80 mw), 640 (75 mw)], PRIME 95B Monochrome Digital Camera and 100x 1.45 NA objective. Objective lenses used included CI Plan Apo Lambda 60x 1.40 NA and Plan Apo Lambda 100x 1.45 NA. Laser power was set to 10% for all channels. Images were acquired using NIS-Elements Advanced Research Software on a Dual Xeon Imaging workstation. RNA/DNA FISH in Figure 4A and Figure S7A-D were deconvoluted using standard parameters. Maximum intensity projection of z-stacks and adjustment of brightness and contrast were performed using Fiji software.

#### Live-cell imaging

For live-cell imaging in Figure 1C, Figure 2C, Figure 2E, Figure 3A, Figure 3C, Figure 4D, Figure 4F, Figure S1C and S1D, Figure S2A, Figure S4C, Figure S4G, Figure S4I, Figure S6A, Figure S6D and Figure S7E, COLO320-DM TetO-EGFP and PC3 TetO-EGFP cells expressing H2B-mCherry and TetR-GFP were seeded into four-well chamber slides (ibidi) 24-72 hours before imaging. Treatments or siRNA transfections were conducted as detailed above 24 hours after seeding. Live-cell imaging was performed at 37 °C heated chamber with humidification and 5 % CO2 using Nikon SoRa Spinning Disk Confocal system, Borealis microadapter, Perfect Focus 4, motorized turret and encoded stage, 5-line laser launch [405 (100 mw), 445 (45 mw), 488 (100 mw), 561 (80 mw), 640 (75 mw)], PRIME 95B Monochrome Digital Camera, and CFI Apo TIRF 60x 1.49 NA objective lens (W.D. 0.12mm). The objective lens used was the CFI Apo TIRF 60x 1.49 NA objective lens (W.D. 0.12mm) in super-resolution mode. Laser powers were set at 5 % power and an exposure time of 600 ms was utilized. Images were acquired using NIS-Elements Advanced Research Software on a Dual Xeon Imaging workstation and denoised using default parameters. Time-lapsed images were taken of mitotic cells every 3-10 minutes beginning at metaphase or anaphase until daughter nuclei were reformed. Maximum intensity projection of z-stacks and processing of images, including the adjustment of brightness and contrast were performed using Fiji software.

#### Image analysis

All image analysis was carried out on Fiji software. To quantify ecDNA nuclear segregation fidelity in Figure 1B, and Figure S1A and S1B, newly formed daughter cells were identified via the presence of Aurora B. Total oncogene FISH signal in the maximum projection of each daughter nucleus was determined. ecDNA/HSR signal was classified as the mean plus one standard deviation. The proportion of nuclear ecDNA signal to total ecDNA signal (nuclear plus cytosolic) was calculated to determine ecDNA nuclear fidelity (Figure 1B). To quantify RNA/DNA FISH signal as seen in Figure S3C and S3D, Figures 4B and 4C, the average signal of each FISH probe was determined from the maximum projection images at ecDNA+ regions in the primary nucleus (Figure S3C and S3D, Figures 4B and 4C) or micronucleus (Figure S3C and S3D). Background signal was subtracted and each RNA FISH probe signal was normalized to ecDNA DNA FISH signal at the region of interest.

For all live-cell imaging, all images were denoised on Nikon NIS-Elements Advanced Research Software and then further processing was carried out using Fiji software, including generation of maximum intensity projections, adjustment of brightness and contrast, and generation of time-lapse movies. Image analysis was carried out using denoised images on NIS-Elements software. To quantify ecDNA cytosolic mis-segregation, individual z-slices were analyzed for H2B and TetR signal outside of the main chromosome mass for each time-point collected. Following the end of telophase, once nuclear envelope reformation and chromosome decondensation began to occur via the H2B channel, any ecDNA signal remaining outside the daughter nucleus, including ecDNA+ micronuclei, was classified as a cytosolic mis-segregation.

#### HiC Library preparation and sequencing

To obtain mitotic samples, COLO320-DM cells were treated with 100 ng/mL nocodazole for 4 hours. Asynchronous samples were left untreated. After four hours, cells were washed once, trypsinized and fixed in 2 % PFA in PBS for 12 minutes. Following incubation, PFA was removed, and cells were washed 2X in PBS.

To prepare cells for sorting, cells were permeabilized (10 mM Tris-HCl pH 8.0, 10 mM NaCl, 0.2 % IGEPAL CA-630, EDTA protease inhibitor in PBS) for 15 minutes at room temperature. Following permeabilization, cells were incubated in blocking buffer (1 % BSA) for 1 hour, followed by MPM2 primary antibody diluted 1:500 in blocking buffer for 1 hour with frequent agitation. Cells were washed twice in wash buffer (0.2 % BSA in PBS), and resuspended in secondary antibody diluted 1:1,000 in blocking buffer for 45 minutes in dark. Samples were then washed with wash buffer 2 times before being resuspended in FACS buffer (0.2 % BSA in PBS) and propidium iodide (100 μg/mL). Cells were sorted using a 130 µm nozzle at 4 °C on a BD FACS Aria (BD Biosciences) at the MSKCC Flow Core Facility. To determine sorting purity, a small fraction of cells were sorted and immediately analyzed on Fortessa. Following sorting, cells were frozen as pellets at −80 °C and further processed for Hi-C at the Epigenetics and Innovation Research Laboratory (ERIL) at MSKCC. Arima-HiC kit streamlined workflow was followed as per manufacturer’s protocol. Arima-Hi-C kit (Arima, A510008) in conjunction with Arima Library prep module. Per each Hi-C condition, approximately 4 × 10^6^ cells were collected, and approximately 200 ng of DNA was used to prepare libraries. Libraries were sequenced using the NovaSeq 6000 flow cell in PE100 mode at Integrated Genomics Operation (IGO) at MSKCC.

#### HiC sequencing analysis

Reads were trimmed of the 5’ end adapter sequences using Cutadapt (version 2.4), aligned to the hg38 genome using bowtie2 (version 2.4.4), and all valid pairs were filtered for using HiC-Pro (version 3.1.0).

All valid pairs were then binned into 1 Mb bins, and the amplicon to linear genome contacts were visualized by filtering for all trans-chromosomal contacts with the bins containing the amplicon region (chr8:127000000-129000000). All bins that are paired with the amplicon region will then be sequentially graphed on a linear chromosome map with the y-axis showing the log 10 fold change of the observed over expected contacts within each bin.

